# Response to “No evidence of functional co-adaptation between clustered microRNAs”

**DOI:** 10.1101/313817

**Authors:** Yirong Wang, Hong Zhang, Jian Lu

**Affiliations:** State Key Laboratory of Protein and Plant Gene Research, Center for Bioinformatics, School of Life Sciences, Peking University, Beijing, 100871, China

**Keywords:** miRNAs, target, functional co-adaptation, drift-draft

## Abstract

microRNAs (miRNAs) are a class of endogenously expressed small non-coding RNAs that regulate target genes at the post-transcriptional level. One significant feature of miRNA is that their genomic locations are often clustered together in the genome. In a previous study (Wang, et al. 2016), we proposed a “functional co-adaptation” model to explain how clustering helps new miRNAs survive and develop functions during long-term evolution. In a manuscript recently posted at bioRxiv (doi:10.1101/274811), Marco claimed that he re-analyzed our data and came to a different conclusion. However, we found his analyses were conducted in an inappropriate approach. He also claimed that the absence of substitution in highly conserved miRNAs does not support the “functional co-adaption” model based on the misunderstanding of our model. In summary, the analyses and claims of Marco, which are flawed, do not refute our model.

---

miRNAs are a class of endogenously expressed small noncoding RNAs (∼22 nt in length) that down-regulate the expression of target genes at the post-transcriptional level. A salient feature is that many animal miRNAs are clustered into discrete genomic regions (Lagos-Quintana et al. 2001; Lau et al. 2001; Lai et al. 2003; Altuvia et al. 2005; J. Graham Ruby et al. 2007; Marco et al. 2013; Mohammed, Siepel, et al. 2014). The clustering patterns suggest that miRNAs in the same cluster might be co-transcribed (Baskerville and Bartel 2005; Saini et al. 2007; Ozsolak et al. 2008; Wang et al. 2009; Ryazansky et al. 2011) and be functionally related by targeting the same gene or different genes in the same biological pathway (Bartel 2004; Grun, et al. 2005; Kim and Nam 2006; Yu, et al. 2006). For example, the *mir-17∼92* cluster plays an important role in mammalian development and tumorigenesis (O’Donnell, et al. 2005; Lu, et al. 2007; Ventura, et al. 2008; Xiao, et al. 2008). Gene deletion experiments suggest members in the *mir-17∼92* cluster have essential and overlapping functions (Ventura, et al. 2008). The *mir-106b∼93∼25* and *mir-222∼221* clusters are upregulated and modulate G1/S phase transition in gastric cancer, and members of the two cluster have functional associations by targeting genes in the Cip/Kip family members of Cdk inhibitors (Kim, et al. 2009). The brain specifically expressed *mir-379∼410* cluster is required for the activity-dependent development of hippocampal neurons, and multiple miRNAs from the cluster are necessary for the correct elaboration of the dendritic tree (Fiore, et al. 2009). miRNAs in *mir-23a∼27a∼24-2* cluster also have cooperative effects in various health and diseased conditions (Chhabra, et al. 2010).

We recently proposed a “functional co-adaptation” model to systematically investigate the functional relatedness of clustered miRNAs (Wang, et al. 2016). We provided several lines of evidence to support the “functional co-adaptation” model. First, we found the observed number of genes co-targeted by miRNAs in the same cluster but with different seeds are significantly higher than the number obtained by random permutations. Second, we found genes targeted by multiple miRNAs from the same clusters, in general, have lower expression levels than genes targeted by multiple miRNAs from distinct clusters. Third, we show that the miRNAs in the same cluster with different seeds tend to target genes in the same biological pathways. Fourth, we transfected four members of the *mir-17∼92* cluster into human 293FT cells individually and quantified the alteration of mRNA abundance with deep-sequencing, which verified the overlapping of target genes experimentally. Fifth, we experimentally determined the target genes of *miR-92a*, the founding member of the *mir-17∼92* in *Drosophila*,and examined the relationship between the target genes of *miR-92a* in *Drosophila* and the target genes of the *mir-17∼92* cluster in humans. Our experimental results well supported the “functional co-adaptation” model. Finally, we also conducted evolutionary analysis to show that positive Darwinian selection drives the evolution of the newly formed miRNA clusters in both primates and *Drosophila*.

In a manuscript recently posted at bioRxiv (Marco, 2018; doi:10.1101/274811), Marco claimed that he re-analyzed our data and found “No evidence of functional co-adaptation between clustered microRNAs”. Marco claimed that the observed overlap of target genes by the clustered miRNAs are mostly caused by the similarity between two seed sequences in the *miR-182/183/96* cluster. Marco argued that clustered miRNAs from different miRNA families do not share more targets than expected by chance after correcting for these factors. Marco also argued that our permutation tests by shuffling miRNA-target inactions would lead to spuriously low *P* values. Moreover, Marco also raised a series of other critiques about the “function co-adaptation” model in different versions of his comments. Unfortunately, Marco’s critiques are based on flawed analysis and misled concept of miRNA evolution, which are summarized as follows.

The major concern Marco raised is whether the observed number of genes targeted by at least two conserved miRNAs with different seeds from the same miRNA clusters is statistically higher than the number obtained under the assumption of randomness. In our previous study (Wang, et al. 2016), to test whether miRNAs in the same clusters tend to regulate overlapping sets of genes, we obtained expression profiles of miRNAs and mRNAs from five tissues of human males as determined in previous study (Brawand, et al. 2011; Meunier, et al. 2013). Since the co-adaption of clustered miRNAs is the result of co-evolution between miRNA and target sites, in the permutation analysis, we first shuffled the co-expressed seed:target pairing (TargetScan *P_CT_* > 0.5), and then we tested how many genes were targeted by at least two miRNAs (with distinct seeds) in the same clusters. These permutation tests were performed for 1,000 replicates. By this way, the conservation level and length of 3’ UTR of target mRNAs, the number of miRNAs for each target gene, and the compositions of each miRNA cluster are fully controlled. Applying this procedure to the pooled dataset of miRNA-mRNA co-expression from different tissues, the result of Wang et al. was successfully reproduced (Fig. 1). When the co-expression data of each tissue was analyzed individually, the similar pattern was still observed (Fig. 1). Importantly, when the *miR-182/183/96* cluster was excluded, we can still observe similar patterns (Fig. 2).

**Figure 1.**
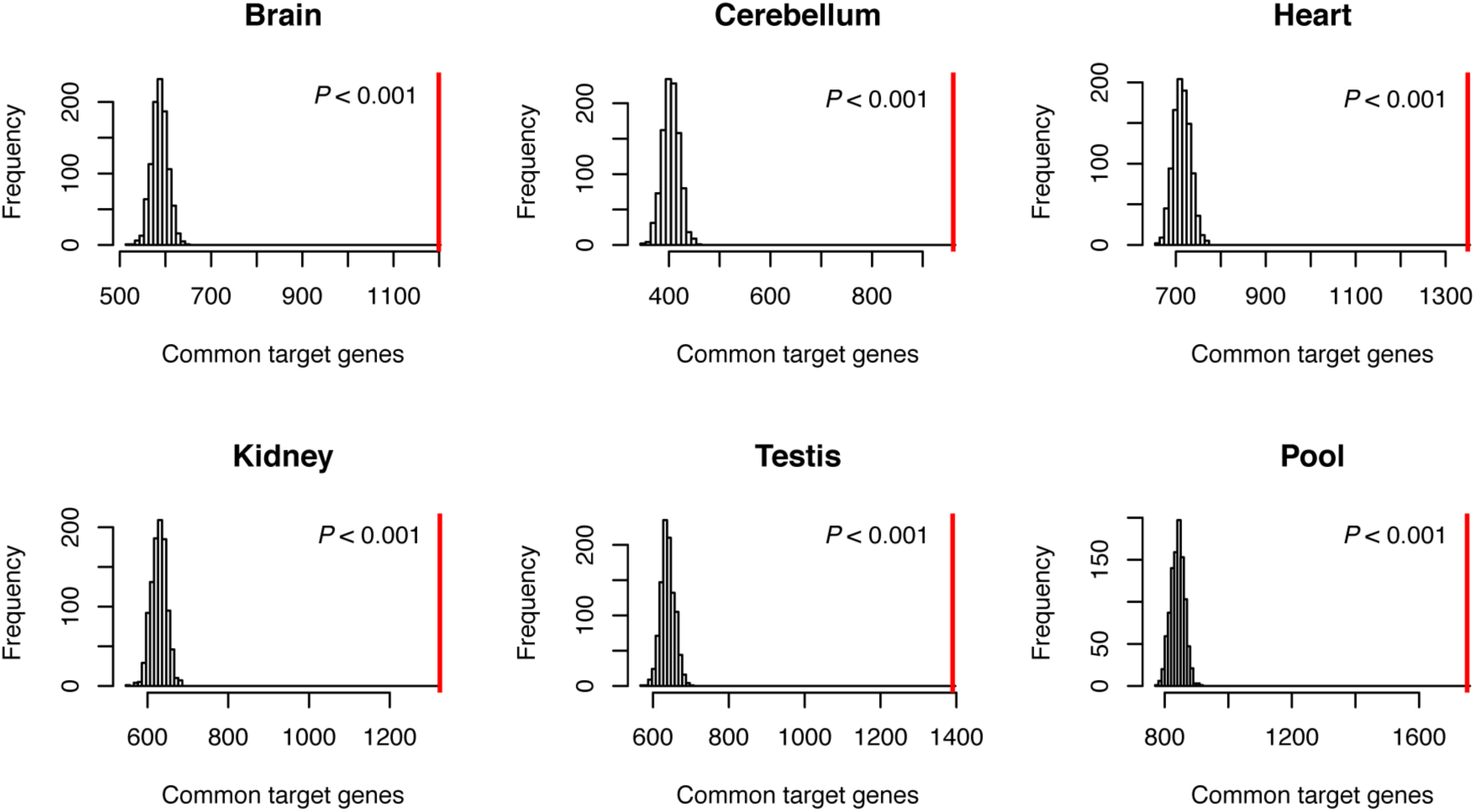
Permutation analysis of common target genes of miRNAs from the same cluster in each tissue or pooled data by shuffling co-expressed seed:target pairs. 1,000 replicates were performed for each panel. The observed number of common targets was indicated with vertical red line and the proportion of simulations yielding a number larger than observed value was shown at top right.

**Figure 2.**
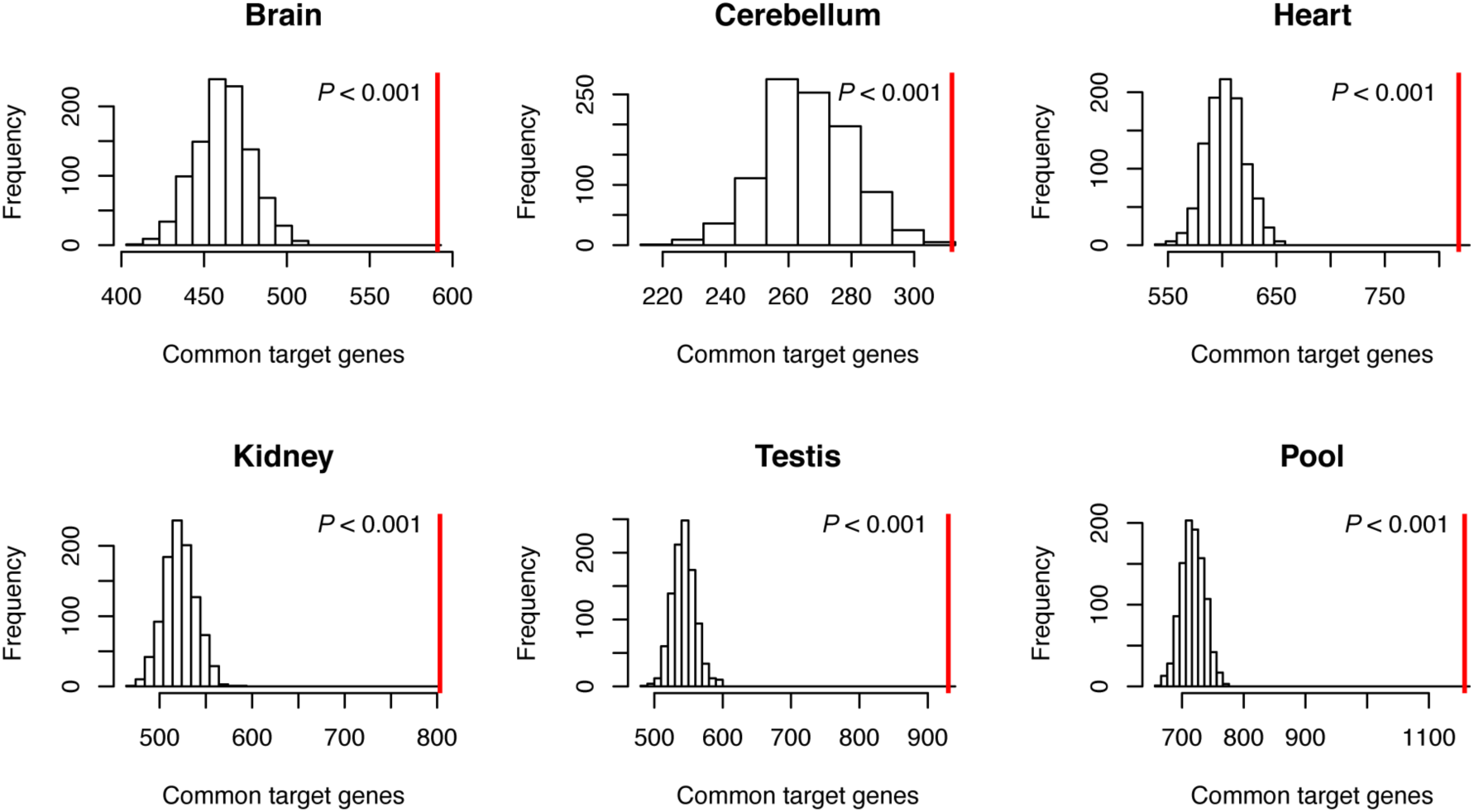
Permutation analysis of common target genes of miRNAs from the same cluster in each tissue or pooled data by shuffling co-expressed seed:target pairs. Similar as Fig. 1 but *miR-182/183/96* cluster was excluded in the analysis.

By contrast, Marco failed to reproduce our results because he only shuffled the location of the miRNAs and kept the seed: targeting pairing unaltered. Marco obtained a pattern that the observed number slightly but still significantly higher than the expected number under the assumption of randomness (*P* = 0.0359). Since the difference between the observed and expected numbers are quite smaller obtained by Marco compared to what we obtained (Wang, et al. 2016), Marco argued that our results are mainly caused by the similarity of the targets between two seed sequences of the *miR-182/183/96* cluster, and “the expected high number of common targets between pairs of microRNAs that have a large number of targets each”. However, Marco’s permutation tests are biased and flawed. Here, the condition to be tested is whether miRNAs with distinct seeds from the same cluster have more common target genes than expected under randomness, rather than to test whether miRNAs are clustered as Marco conducted.

Despite the numerous exchanges between Marco and us on the detailed permutation test procedures, Marco consistently argued that our permutation tests by shuffling miRNA-target interactions would lead to spuriously low *P* values. To show this point, Marco introduced a hypothetical regulatory network, in which two clustered miRNAs and an additional miRNA target the same gene “a” and each miRNA has *n* additional targets (see Figure 2 of Marco, 2018; doi:10.1101/274811 for details). Marco derived the formula for calculating the probability that a permutation reports a common target between two clustered microRNAs under randomness as

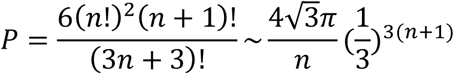

based on “standard combinatorics” (Marco, 2018; doi:10.1101/274811).

Marco argued that based on this formula, one can obtain very small *P* values even when there is no enrichment in common genes targeted by clustered microRNAs. Unfortunately, after careful examination, we found the formula Marco derived is incorrect. The correct formula for *P* value should be:

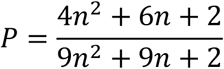

based on basic combinatorics (see Appendix for details).

In Figure 3, we show that the *P* values calculated by Marco are consistently smaller than the *P* values calculated by the formula we derived. To further support our argument, we also followed the scheme presented in Figure 2 of Marco, 2018 (doi:10.1101/274811) and estimated the empirical *P* value by randomly permutating the miRNA:target pairing for 1,000 times at a given *n*. As we show in Figure 3, The *P* values calculated with the correct formula (red curve) are highly consistent with the empirical *P* values, while the *P* values calculated with Marco’s formula are consistently much smaller than the *P* values obtained by simulations. Therefore, Marco’s argument that our permutation tests by shuffling miRNA-target interactions would lead to spuriously low *P* values is based on his erroneous formula in calculating the *P* value. In fact, the correct *P* value formula indicates shuffling miRNA-target interactions would not generate a significant *P* value when there is no enrichment in common genes targeted by clustered microRNAs.

**Figure 3.**
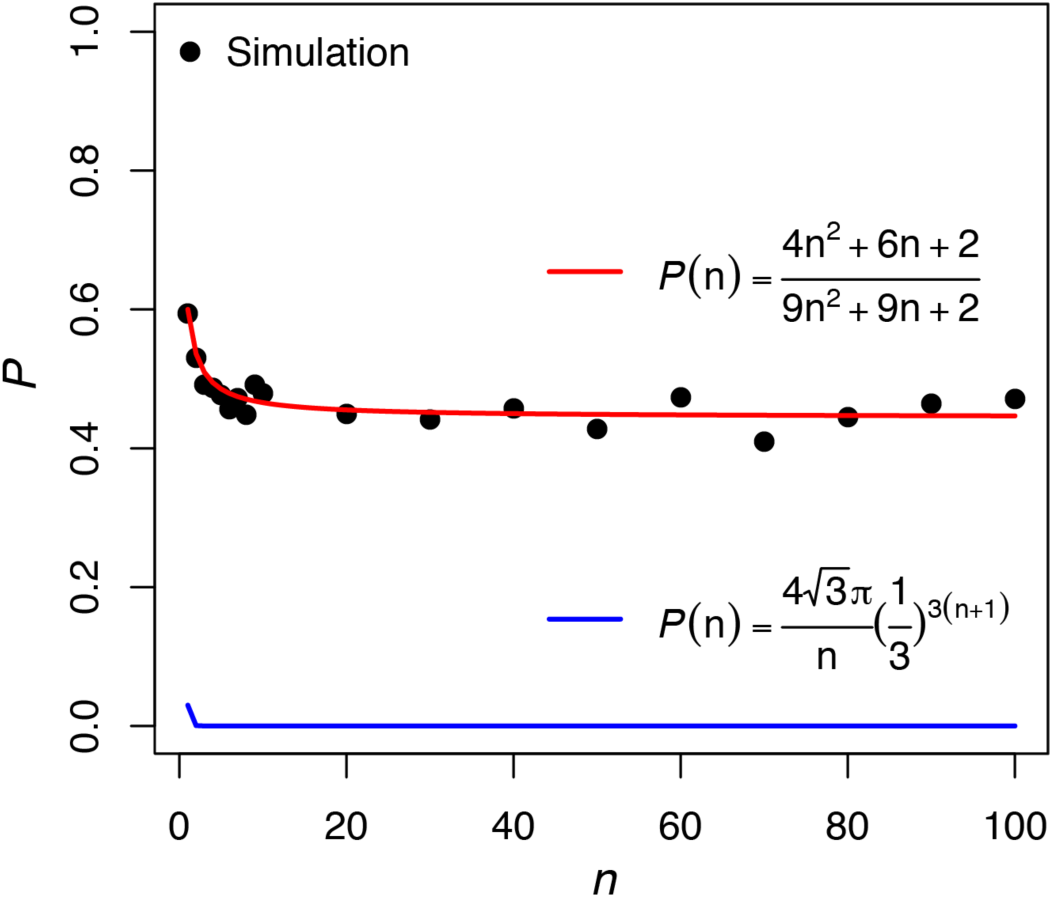
The comparison between the *P* values (y axis) obtained by Marco’s formula (blue line) and the formula derived by us (red line) and the simulation results (black points). In a hypothetical regulatory network, two clustered miRNAs and an additional miRNA target the same gene “a” and each miRNA has *n* (*x-axis*) additional targets, and the *P* value is the probability that a permutation reports a common target between two clustered microRNAs under randomness (see Figure 2 of Marco, 2018; doi:10.1101/274811 for details). The *P* value calculated with Marco’s formula are consistently smaller (< 0.05) even when there is no enrichment in common genes targeted by clustered microRNAs. However, the *P* values calculated with the correct formula (red curve) indicates shuffling miRNA-target interactions would not generate a significant *P* value when there is no enrichment in common genes targeted by clustered microRNAs. The black points are the empirical *P* value obtained by randomly permutating the miRNA:target pairing for 1,000 times at a given *n*. Note our formula (red line) and simulation results (black points) are highly consistent.

Marco also argued that the differences between clustered and non-clustered miRNAs are not significant when he only focused on the pairs of miRNAs that have fewer than 6 common nucleotides at seed regions in a cluster (the Fig. 1C of (Marco 2018)). He argued that the observed overlap between targets in some clustered miRNAs is actually the random consequence of the similarity between their seed sequences, and is not associated with whether the miRNAs are clustered or not. An inspection of his code indicates his critique is based on incorrect calculation of “the fraction of common targets” for a pair of miRNAs. For example, let us suppose that miRA has *x* targets, miRB has *y* targets, and *z* targets are shared between miRA and miRB. Marco calculated the fraction of common targets as *z*/min(*x,y*) in his Fig. 1C, while calculated the fraction of common targets as *z*(*x+y*)/(2*xy*) in his Fig. 1D. Obviously, using the harmonic mean (*z*(*x+y*)/(2*xy*)) is more appropriate to calculate the fraction of common targets, as employed by Marco in his Fig. 1D. Interestingly, when we repeated his analysis in Fig. 1C with the harmonic mean (*z*(*x+y*)/(2*xy*)) approach as he did in Fig. 1D, we found a pair of miRNAs with seed similarity less than 6 nucleotides have a significantly higher fraction of common targets if the two paired miRNAs are from the same miRNA cluster (*P* = 0.008, Fig. 4). Therefore, the results in the Fig. 1C of (Marco 2018) are based on incorrect analytical procedures.

**Figure 4.**
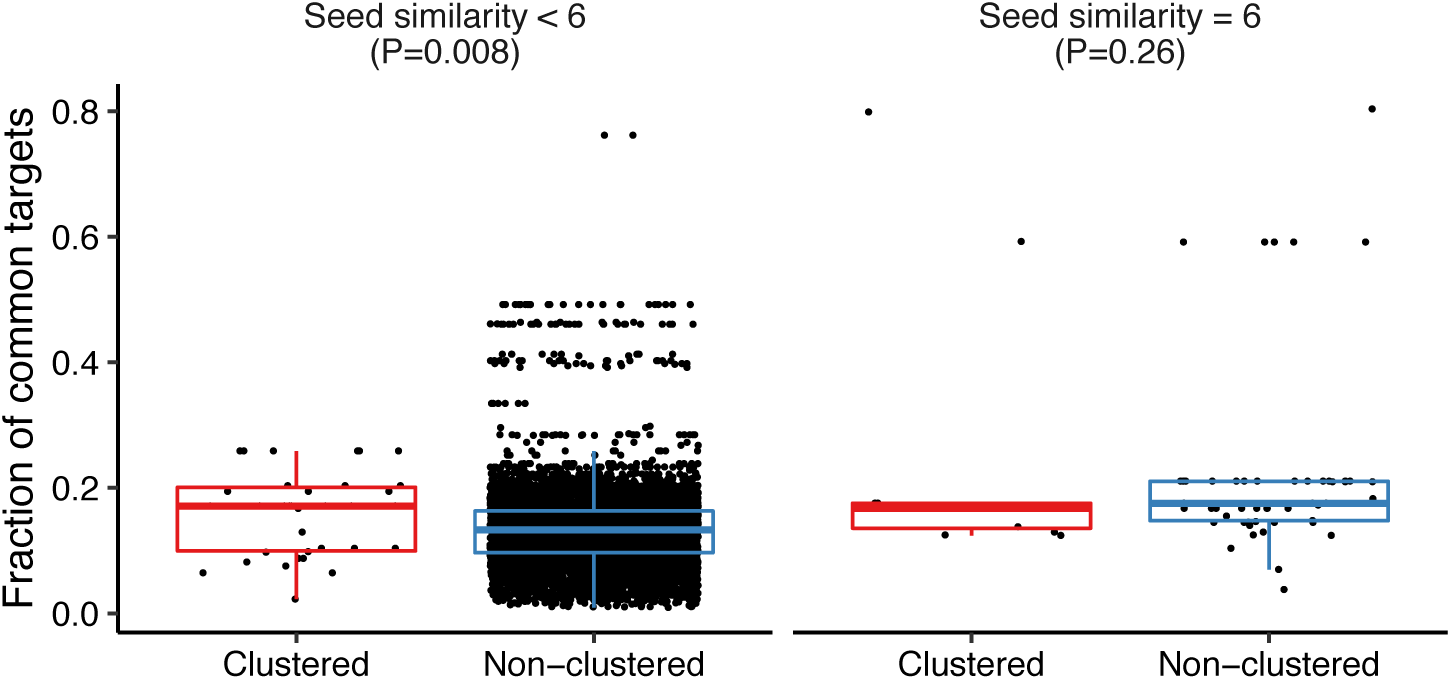
The distribution of the fraction of common targets for a pair of miRNAs that are from the same cluster (Clustered) or not (Non-clustered). Paired miRNAs with a seed similarity less than or equal to six nucleotides were compared separately with the Wilcoxon rank-sum tests.

Notably, both our original analysis (Wang, et al. 2016) and Marco (2018) only considered the evolutionarily conserved target sites of the evolutionarily conserved miRNAs (only miRNAs that have target sites with TargetScan *P_CT_* > 0.5 were used in the permutation analyses). Here we considered all the miRNAs that have at least one target site (not necessarily conserved) with weighted context++ score (WCC) < −0.3 (Grimson, et al. 2007; Agarwal, et al. 2015). Target sites with WCC < −0.3 are usually located in optimized genomic context for efficient repression of target mRNAs. With the criteria of WCC < −0.3, only 125 (6%) of 2,081 common targets of clustered miRNAs are contributed by *miR-182/183/96*, which would remarkably reduce the potential bias caused by *miR-182/183/96*. We consistently observed that clustered miRNAs have significantly more common target genes than expected before (empirical *P* = 0.0037, Fig. 5A) or after (empirical *P* = 0.0088, Fig. 5B) excluding *miR-182/183/96* cluster in the permutation tests by shuffling miRNA loci as Marco performed. A more significant trend was observed when the permutation tests were performed by shuffling co-expressed seed: target pairs (*P* < 0.0001 in both Fig. 5C and 5D) as conducted in our original study (Wang, et al. 2016). Therefore, we reaffirmed the thesis that clustered miRNAs have more common target genes than expected using miRNA targets with WCC < −0.3.

**Figure 5.**
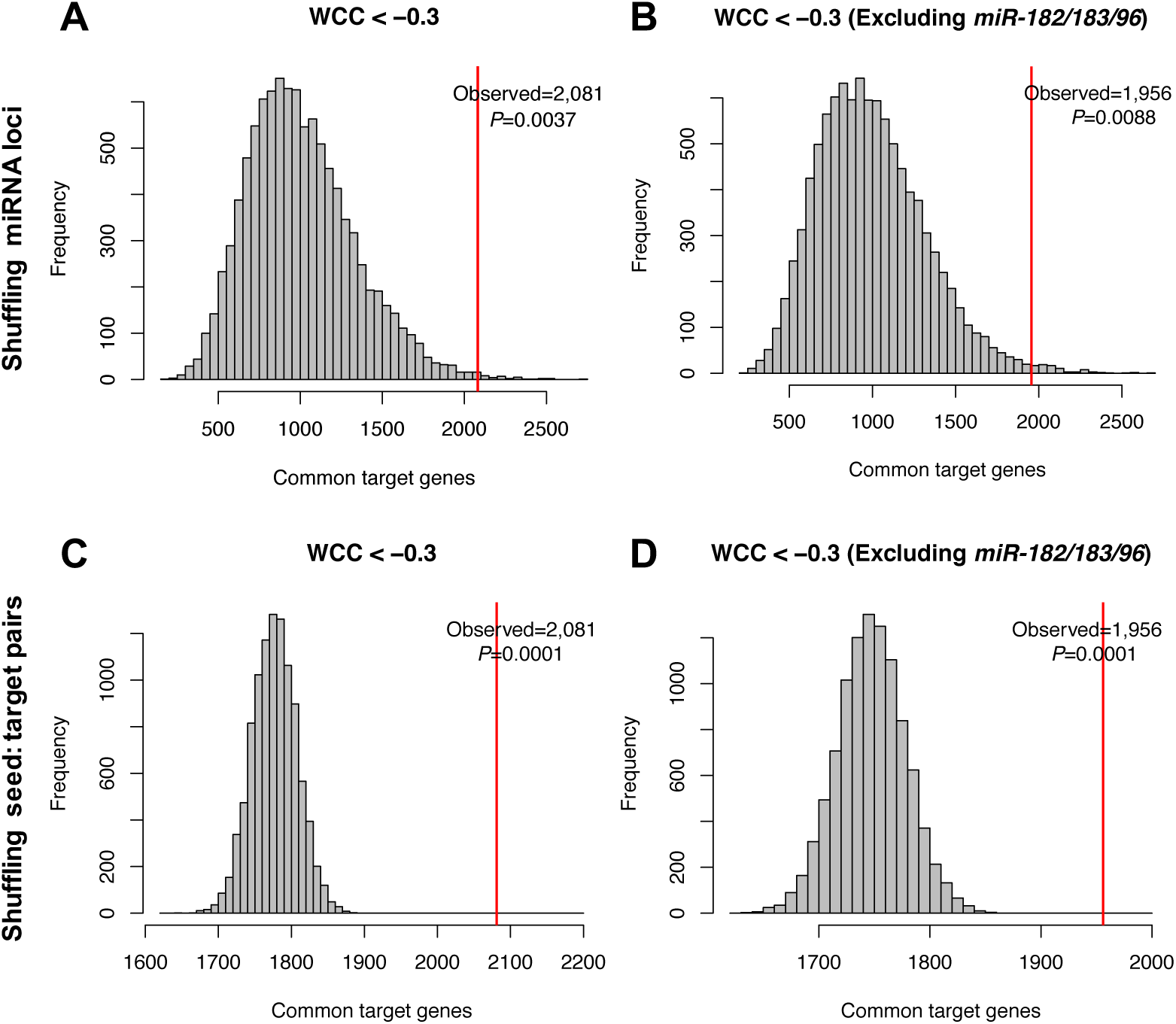
Permutation analysis of common target genes of clustered miRNAs. (**A** and **B**) Permutation analysis by shuffling miRNA loci using miRNA target genes with weighted context score++ (WCC) < −0.3 before (A) or after (B) excluding *miR-182/183/96* cluster. (**C** and **D**) Permutation analysis by shuffling co-expressed seed: target pairs using miRNA target genes with WCC < −0.3 before (C) or after (D) excluding *miR-182/183/96* cluster. All the human miRNAs and their targets with WCC < −0.3 were analyzed. Only co-expressed miRNA:target pairs in human tissues were considered. 10,000 replicates were performed for each permutation test. The observed numbers of common targets were indicated with red lines. The empirical *P* value was calculated as the fraction of simulations yielding a number of common targets larger than the observed one.

Furthermore, it is hard to understand why Marco argued that the targets shared between *miR-183* and *miR-96* in the *miR-182/183/96* cluster should be excluded from the analysis. The seeds of *miR-183-5p* and *miR-96-5p* are very similar: AUGGCAC and UUGGCAC for the former and latter, respectively. However, BLAST2SEQ analysis between the precursor sequences of human *mir-183* and *mir-96* does not find significant similarity, suggesting these two miRNA precursors are unlikely to be duplicated miRNAs. Instead, the functional co-adaptation model might well explain the large number of target genes shared between these two miRNAs: During long time evolution, the adaptive changes in miRNA seed region or target sites on mRNAs drive the clustered miRNAs to regulate the same or functionally related genes. Therefore, this cluster serves as a strong evidence that convergent evolution has occurred between the seeds of *miR-183* and *miR-96* due to functional coadaptation.

Curiously, Marco did not report his re-analysis results of the *miR-17∼92* cluster over-expression data we generated (Wang, et al. 2016). Many previous studies have demonstrated that the *mir-17∼92* cluster plays an important role in tumorigenesis, development of lungs and immune systems (O’Donnell, et al. 2005; Lu, et al. 2007; Ventura, et al. 2008; Xiao, et al. 2008), and deletion of the *mir-17∼92* cluster revealed that miRNAs in this cluster have essential and overlapping functions (Ventura, et al. 2008). Furthermore, we found the conserved target genes shared between members of the *mir-17∼92* cluster is significantly higher than the simulated ones (Fig. 6). Importantly, we selected four distinct mature miRNAs in the *miR-17∼92* cluster (*miR-17, miR-18a, miR-19a*, and *miR-92a*) and transfected each miRNA mimic as well as the miRNA mimic Negative Control (NC) into human 293FT cells (Wang, et al. 2016). With high-throughput mRNA-Seq, we found the predicted target genes (TargetScan *P_CT_* > 0.5) of each transfected miRNA are significantly more down-regulated than genes that do not have the target sites (Figure 4 of Wang et al. 2016). We identified 301, 55, 345 and 268 high-confidence target genes (TargetScan *P_CT_* > 0.5) for *miR-17, 18a, 19a* and *92a* respectively that were down-regulated with log2(FoldChange) < −0.1 in the corresponding miRNA transfection experiments (totally 775 high-confidence genes after removing overlapping genes, Figure 4I of Wang et al. 2016). Among these 775 high-confidence target genes, 172 were targeted by at least two out of the four miRNAs, significantly higher than the number obtained by randomness (*P* < 0.001, see Figure 4I and Table S8 of Wang et al. 2016 for details). These results well support the “functional co-adaptation model” we proposed. If there is really “No evidence of functional co-adaptation between clustered microRNAs” as Marco argued, how can one explain these observed patterns?

**Figure 6.**
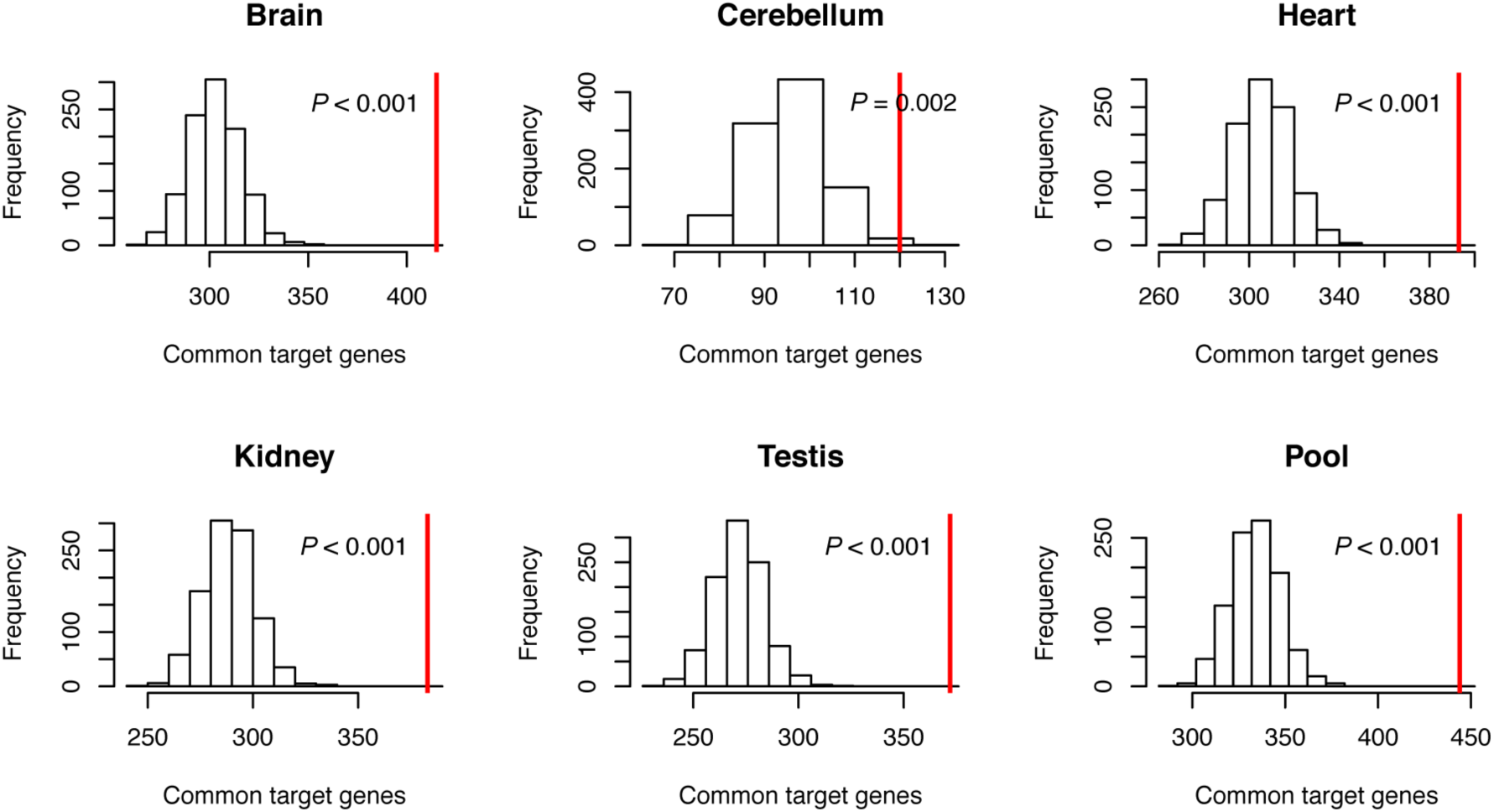
The number of observed (red) and simulated common target genes of conserved miRNAs in *mir-17∼92* cluster in the permutation analysis of Fig. 1.

Moreover, Marco did not fully understand the “functional co-adaption” model, which led him to make the argument that “microRNAs in a cluster are primarily under positive selection” (April/20/2018 version of doi:10.1101/274811). Although our population genetic analysis suggests Darwinian selection drives the evolution of the newly formed miRNA clusters in primates and in *Drosophila*, our model does not necessarily suggest all the clustered miRNAs are driven by positive selection (Wang, et al. 2016). What we proposed is that, new miRNAs originating nearby a pre-existed miRNA would have a higher chance to be maintained in the initial stage of cluster formation due to the tight genetic linkage. Then positive Darwinian selection might drive the newly emerged miRNAs to develop functions related to the pre-existing miRNAs in the same cluster or drive the evolution of all the new miRNAs in the same cluster to develop related functions during the long-term evolution. Once the cluster is fully established, the miRNAs in the same cluster will be maintained by purifying selection and become highly conserved after that (Wang, et al. 2016). Thus, one could not expect to observe signature of ongoing positive selection in the well-established clusters such as the *miR-17∼92* or the *miR-182/183/96* cluster which are ancient and conserved after the establishment, as Marco did. Marco’s observation that “both seed sequences (of *miR-183-5p* and *miR-96-5p*) have been conserved since their origin and, therefore, there is no evidence of substitutions happening in the seed of these microRNAs for the last 600 million years” could not refute our model. Marco also used the deep conservation of the clustered miRNAs in other clusters (*mir-106b∼25* cluster, *mir-23b∼24* cluster, and *mir-379∼410* cluster of Fig. S1 in the April/20/2018 version of his manuscript) to argue against the “functional co-adaptation” model. The deep conservation of the seed sequences as Marco showed can only suggest these miRNAs are conserved due to extremely strong selective constraints during vertebrate evolution. These observations do not provide any evidence to defy the “functional co-adaptation” model since one cannot tell whether the miRNAs have changed since emergence as no outgroup sequence available.

Based on the observation that *Drosophila* new miRNAs often arose around the pre-existing ones to form clusters, Marco and colleagues proposed a “drift-draft” model which suggests that the evolution of miRNA clusters was influenced by tight genetic linkage and largely non-adaptive (Marco, et al. 2013). Under such a model, the motifs of the pre-existing miRNAs would protect new miRNAs to be transcribed and processed properly since those motifs were already interacting with the miRNA processing machinery. Thus, the *de novo* formed new miRNAs are sheltered by the established ones in the same cluster because mutations that abolish the transcription or processing of the new miRNA will affect the pre-existing ones as well and are hence selected against. On the other hand, if a *de novo* formed miRNA is located in a discrete locus, it will have a higher probability to degenerate, either by mutations abolishing its transcription or by mutations impairing its processing. Although Marco argued the “drift-draft” and “functional co-adaptation” models are mutually exclusive, we did not think the “functional co-adaptation” we proposed is strictly “an alternative to the drift-draft model”.

Our previous results and others suggest that many newly-emerged miRNAs are evolutionarily transient, with a high birth-and-death rate (Berezikov et al. 2006; Rajagopalan et al. 2006; Lu, Shen, et al. 2008; Lu et al. 2010). Therefore, it is possible that the newly emerged miRNAs in the clusters would be sheltered by the pre-existing established miRNAs. However, the protection effect alone cannot explain why miRNAs in the same cluster have significantly higher numbers of overlapping target genes. Moreover, many *de novo* formed novel miRNAs will degenerate even after they are fixed in the populations if they are not maintained by functional constraints (Berezikov et al. 2006; Lu, Shen, et al. 2008). Thus developing functions related to the pre-existing miRNAs will help the novel miRNAs to survive and stabilize. The “functional co-adaptation” model we proposed well accounts for the evolution and function of *de novo* formed new miRNAs in the clusters (Figure 2D of Wang et al. 2016). Since miRNAs in the same clusters are usually co-transcribed temporally or spatially (see below for details), the newly formed miRNAs might gradually develop functions to target genes that are related to the pre-existing miRNAs in the same cluster; or multiple *de novo* formed new miRNAs in the same cluster interplay to regulate overlapping sets of target genes. Therefore, although miRNAs in the same cluster have independent origins, they might regulate overlapping sets of target genes through convergent evolution. After that, the clustering patterns of miRNAs and the modular regulation of target genes will be stabilized by natural selection during long-term evolution. Of course, the evolutionary process of miRNAs is also companied by the co-evolution of the target sites, which might also potentially affect the base compositions or even the length of a 3’ UTR. In a separate study, we also showed the target sites of miRNAs also experienced frequent births and deaths (Luo, et al. 2018). But the evolution of the target sites alone would not cause the clustering pattern of miRNAs. A plausible scenario is that after a new miRNA originates in a cluster, the substitutions that change the sequences and expression of the new miRNAs, the interactions between miRNAs in the same cluster, and the co-evolution between miRNAs and the target sites or the 3’ UTRs, jointly affect the evolution of the clustering pattern of miRNAs.

Understanding the molecular mechanisms and evolutionary principles of the miRNA clustering would deepen our understanding of the regulatory roles of miRNAs in various biological processes or diseases. The “functional co-adaptation” model we propose is well supported by evolutionary and functional genomic data.

## Supporting information

Supplementary codes

## Appendix

In a hypothetical regulatory network where two clustered miRNAs (miR-A and miR-B) and an additional miRNA (miR-C) target the same gene “a” and each miRNA has *n* additional targets (see Figure 2 of Marco, 2018 doi:10.1101/274811 for detailed description), the total number of all possible pairing between the three miRNAs and 3*n* + 1 targets is:

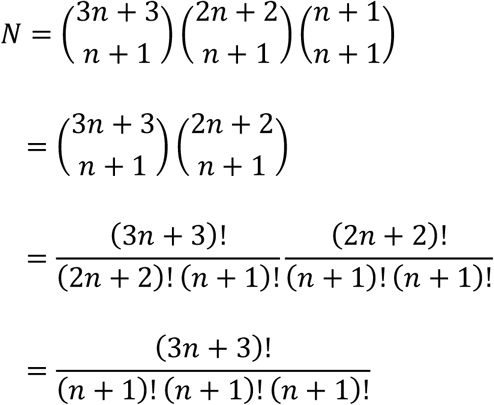

“a” is the only gene that could be targeted by more than one miRNA since other targets only interact with a single miRNA. Since “a” has three interactions, there are three possible scenarios where both the clustered miR-A and miR-B could target gene “a”: (1) two interactions of gene “a” are taken by miR-A, and the remaining one is taken by miR-B; (2) one interaction of gene “a” is taken by miR-A, and the remaining 2 are taken by miR-B; (3) Both miR-A and miR-B takes one interaction of gene “a”, and the remaining one is taken by “miR-C”. We can obtain the number of possible combinations under each of the three scenarios as:

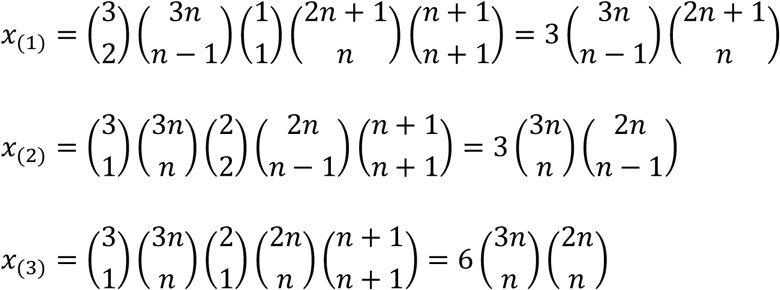

The total number of combinations where gene “a” is targeted by both miR-A and miR-B is:

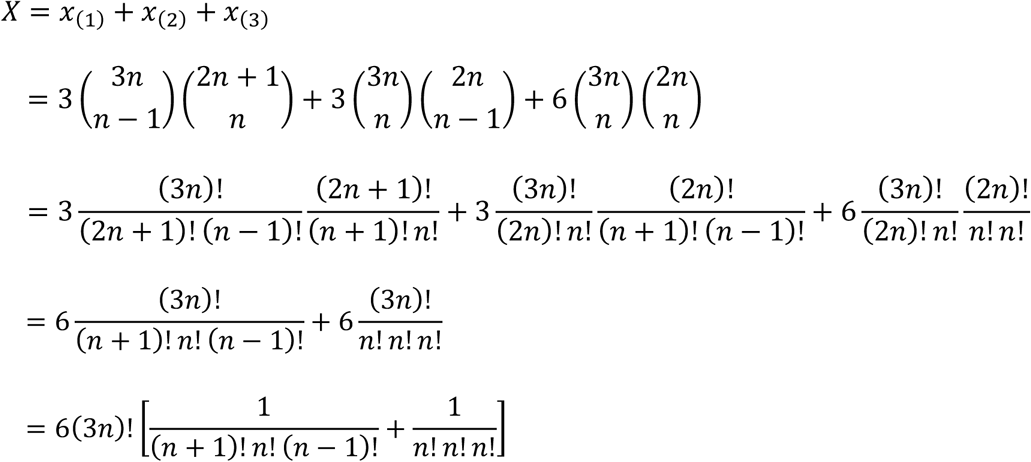

Therefore, the probability that both miR-A and miR-B target gene “a” is:

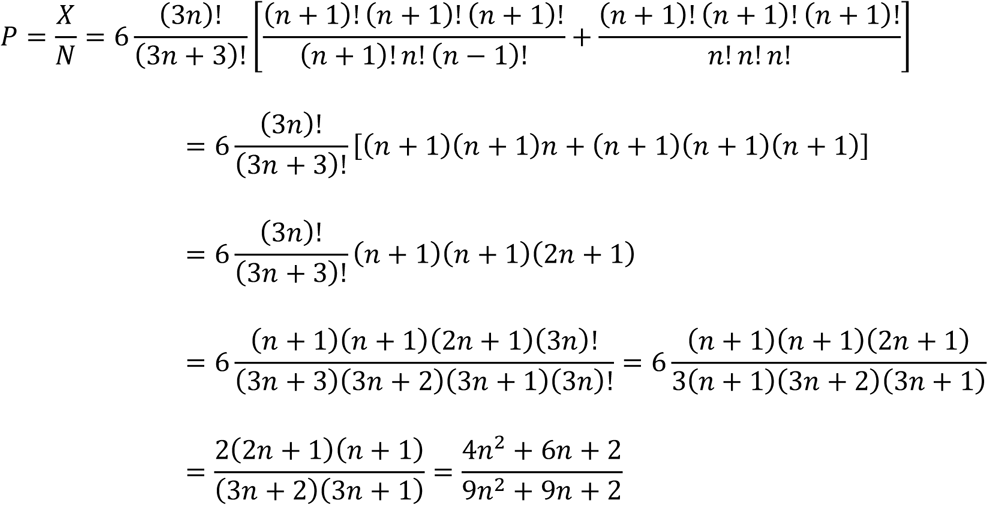

