## Supplementary codes for "Response to “No evidence of functional co-adaptation between clustered microRNAs”": Figure1_Figure2_Figure6.Rmd

````
---
title: "R Notebook"
output: html_notebook
---

### Load datasets
```{r}
library(data.table)

## prepare data ##################################################################################
## refseq 2 ens
ref2ens <- fread('zcat data/ensembl_refseq_protein_id.parsed.gz')
setnames(ref2ens, names(ref2ens), c("ens","tr","refseq","aa"))

## mRNA expression
mrna <- fread("zcat data/henrick_mRNA_expression.txt.gz", header = FALSE)
setnames(mrna, c("ens","exp","tissue"))

mrna.clean <- mrna[ref2ens, on = 'ens', nomatch = 0, allow.cartesian = TRUE]

# mirna cluster
mirna <- fread("data/miRNA_conserved_cluster_seed.txt")
setnames(mirna, names(mirna), c("miR","cluster","seed"))
mirna.uniq <- unique(mirna[,c(2,3), with = FALSE])

## target
target.con <- fread("zcat data/targetscan_site_summary_2015_04backup.txt.gz", header = T)
setnames(target.con, make.names(names(target.con)))
target.con <- target.con[Aggregate.PCT >= 0.5]
target.con <- target.con[, .(refseq = Transcript.ID, seed = miRNA.family)]


# miRNA expression
miRNA <- read.table("data/henrick_miRNA_expression_first19nt.txt",as.is=T)
miRNA2 <- miRNA[miRNA[,7] >=50,][,c(1,6,7,8)]
colnames(miRNA2) <- c("mir","seed","mir_exp","mir_tissue")
setDT(miRNA2)

## observed counts ###############################################################################
## target uniq seed
final.mirna <- mirna.uniq[miRNA2, on = 'seed', nomatch = 0, allow.cartesian = TRUE]
final.mirna.no183 <- final.mirna[cluster != 'chr7_129770405_129774988_-']

target.uniq <- unique(target.con)
#target.uniq[, seed := sample(seed, .N)]

final.tar <- target.uniq[mrna.clean[, .(refseq, ens, tissue)],
                         allow.cartesian = TRUE, nomatch = 0, on = 'refseq']
final.df <- final.tar[final.mirna, on = c('seed', 'tissue' = 'mir_tissue'),
                      allow.cartesian = TRUE, nomatch = 0]
final.smy <- final.df[, .(N = uniqueN(seed)), by = .(cluster, ens)]
uniqueN(final.smy[N > 1, ens])

observed.df <- final.df
remove(final.df)
```

### Shuffling miRNA:target pairs
```{r eval=FALSE, include=TRUE}
if(FALSE) local({
    small.n <- 1000
    require(parallel)

    ## shuffle site position
    set.seed(123)  # note: exact results is not reproducible when `mclapply` is used even with set.seed
    site.shuffle.tissue <- lapply(unique(observed.df$tissue), function(tissue.now){
        res <- mclapply(seq_len(small.n), function(i){
            dtt <- observed.df[tissue == tissue.now]
            dtt <- unique(dtt[, .(seed, ens)])
            x <- sample.int(nrow(dtt))
            dtt[, seed := seed[x]]
            dtt <- dtt[mirna.uniq, on = "seed", allow.cartesian = T, nomatch = 0]
            res <- dtt[, .(N = uniqueN(seed)), by = .(cluster, ens)]
            # all tars
            x1 <- res[N >= 2, uniqueN(ens)]
            # mir17~92
            x2 <- res[cluster == 'chr13_91350617_91351382_+' & N >= 2, uniqueN(ens)]
            c(x1, x2)
        }, mc.cores = 20)
        rbindlist(lapply(res, as.list))
    })
    names(site.shuffle.tissue) <- unique(observed.df$tissue)

    set.seed(123)
    site.shuffle.pool <- mclapply(seq_len(small.n), function(i){
        dtt <- unique(observed.df[, .(seed, ens)])
        x <- sample.int(nrow(dtt))
        dtt[, seed := seed[x]]
        dtt <- dtt[mirna.uniq, on = "seed", allow.cartesian = T, nomatch = 0]
        res <- dtt[, .(N = uniqueN(seed)), by = .(cluster, ens)]
        # all tars
        x1 <- res[N >= 2, uniqueN(ens)]
        # mir17~92
        x2 <- res[cluster == 'chr13_91350617_91351382_+' & N >= 2, uniqueN(ens)]
        c(x1, x2)
    }, mc.cores = 20)
    site.shuffle.pool <- rbindlist(lapply(site.shuffle.pool, as.list))

    ## shuffle site position after removing mir183~182
    set.seed(123)
    site.shuffle.tissue.no183 <- lapply(unique(observed.df$tissue), function(tissue.now){
        res <- mclapply(seq_len(small.n), function(i){
            dtt <- observed.df[tissue == tissue.now]
            dtt <- unique(dtt[, .(seed, ens)])
            x <- sample.int(nrow(dtt))
            dtt[, seed := seed[x]]
            dtt <- dtt[mirna.uniq[cluster != 'chr7_129770405_129774988_-'],
                       on = "seed", allow.cartesian = T, nomatch = 0]
            res <- dtt[, .(N = uniqueN(seed)), by = .(cluster, ens)]
            # all tars
            x1 <- res[N >= 2, uniqueN(ens)]
            # mir17~92
            x2 <- res[cluster == 'chr13_91350617_91351382_+' & N >= 2, uniqueN(ens)]
            c(x1, x2)
        }, mc.cores = 20)
        rbindlist(lapply(res, as.list))
    })
    names(site.shuffle.tissue.no183) <- unique(observed.df$tissue)

    set.seed(123)
    site.shuffle.pool.no183 <- mclapply(seq_len(small.n), function(i){
        dtt <- unique(observed.df[, .(seed, ens)])
        x <- sample.int(nrow(dtt))
        dtt[, seed := seed[x]]
        dtt <- dtt[mirna.uniq[cluster != 'chr7_129770405_129774988_-'],
                   on = "seed", allow.cartesian = T, nomatch = 0]
        res <- dtt[, .(N = uniqueN(seed)), by = .(cluster, ens)]
        # all tars
        x1 <- res[N >= 2, uniqueN(ens)]
        # mir17~92
        x2 <- res[cluster == 'chr13_91350617_91351382_+' & N >= 2, uniqueN(ens)]
        c(x1, x2)
    }, mc.cores = 20)
    site.shuffle.pool.no183 <- rbindlist(lapply(site.shuffle.pool.no183, as.list))

    res <- list(
        site.shuffle.pool = site.shuffle.pool,
        site.shuffle.pool.no183 = site.shuffle.pool.no183,
        site.shuffle.tissue = site.shuffle.tissue,
        site.shuffle.tissue.no183 = site.shuffle.tissue.no183
    )
    saveRDS(res, file = 'results/simulation_shuffle_site_0801.rds')
})
```


### Generate plots
```{r}
capwords <- function(s, strict = FALSE) {
    cap <- function(s) paste(toupper(substring(s, 1, 1)),
                             {s <- substring(s, 2); if(strict) tolower(s) else s},
                             sep = "", collapse = " " )
    sapply(strsplit(s, split = " "), cap, USE.NAMES = !is.null(names(s)))
}

plt.obs <- sapply(unique(observed.df$tissue), function(x){
    observed.df[tissue == x][, .(N = uniqueN(seed)), by = .(cluster, ens)][N > 1, uniqueN(ens)]
})
names(plt.obs) <- unique(observed.df$tissue)
plt.obs <- c(plt.obs, Pool = 1751)

simu.n <- 1000

site.shuffle.server <- readRDS('results/simulation_shuffle_site_0801.rds')

# overall distribution
plt <- c(site.shuffle.server$site.shuffle.tissue,
         list(Pool = site.shuffle.server$site.shuffle.pool))

pdf('plots/Figure1_shuffle_site.pdf', width = 7, height = 4)
op <- par(mar = c(5, 5, 4, 1), mfrow = c(2, 3))
tmp <- mapply(function(x){
    dtf <- as.data.frame(plt[[x]])
    dtf <- head(dtf, simu.n)
    y <- dtf[, 1]
    hist(y,
         seq(min(y) - 10, max(y, plt.obs[x]) + 10, 10),
         xlab = 'Common target genes',
         main = capwords(x))
    abline(v = plt.obs[x], col = 2, lwd = 2)
    p <- sum(y > plt.obs[x]) / length(y)
    legend('topright', legend = p, bty = 'n')
}, names(plt))
par(op)
dev.off()


# remove mir183 ###########################################################
obs.no183 <- sapply(unique(observed.df$tissue), function(x){
    observed.df[tissue == x & cluster != 'chr7_129770405_129774988_-'][
        , .(N = uniqueN(seed)), by = .(cluster, ens)][N > 1, uniqueN(ens)]
})
names(obs.no183) <- unique(observed.df$tissue)
obs.no183 <- c(obs.no183, Pool = 1158)

plt <- c(site.shuffle.server$site.shuffle.tissue.no183,
         list(Pool = site.shuffle.server$site.shuffle.pool.no183))

pdf('plots/Figure2_shuffle_site_no183.pdf', width = 7, height = 4)
op <- par(mar = c(5, 5, 4, 1), mfrow = c(2, 3))
tmp <- mapply(function(x){
    dtf <- as.data.frame(plt[[x]])
    dtf <- head(dtf, simu.n)
    y <- dtf[, 1]
    hist(y,
         seq(min(y) - 10, max(y, obs.no183[x]) + 10, 10),
         xlab = 'Common target genes',
         main = capwords(x))
    abline(v = obs.no183[x], col = 2, lwd = 2)
    p <- sum(y > obs.no183[x]) / length(y)
    legend('topright', legend = p, bty = 'n')
}, names(plt))
par(op)
dev.off()


# mir17-92 #################################################################
obs.mir17 <- sapply(unique(observed.df$tissue), function(x){
    observed.df[tissue == x & cluster == 'chr13_91350617_91351382_+'][
        , .(N = uniqueN(seed)), by = .(cluster, ens)][N > 1, uniqueN(ens)]
})
names(obs.mir17) <- unique(observed.df$tissue)
obs.mir17 <- c(obs.mir17, Pool = 444)

plt <- c(site.shuffle.server$site.shuffle.tissue,
         list(Pool = site.shuffle.server$site.shuffle.pool))

pdf('plots/Figure6_shuffle_site_mir17_92.pdf', width = 7, height = 4)
op <- par(mar = c(5, 5, 4, 1), mfrow = c(2, 3))
tmp <- mapply(function(x){
    dtf <- as.data.frame(plt[[x]])
    dtf <- head(dtf, simu.n)
    y <- dtf[, 2]
    hist(y,
         seq(min(y) - 10, max(y, obs.mir17[x]) + 10, 10),
         xlab = 'Common target genes',
         main = capwords(x))
    abline(v = obs.mir17[x], col = 2, lwd = 2)
    p <- sum(y > obs.mir17[x]) / length(y)
    legend('topright', legend = p, bty = 'n')
}, names(plt))
par(op)
dev.off()
```
````
