## Supplementary codes for "Response to “No evidence of functional co-adaptation between clustered microRNAs”": Figure3.Rmd

````
---
title: ""
output: html_notebook
---

###  Figure 3. P values that both clustered miRNAs miR-A and miR-B target the same gene “a” in a hypothetical regulatory network, where two clustered miRNAs (miR-A and miR-B) and an additional miRNA (miR-C) target the same gene “a” and each miRNA has n additional targets.

#### Simulation
```{r}
library(data.table)

simu <- function(n){
    miRs <- c('A', 'B', 'C')
    tars <- c(0L, seq_len(n * 3))
    
    dtt <- data.table(
        miR = c(miRs, rep(miRs, each = n)),
        tar = rep(tars, c(3, rep(1, n * 3)))
    )
    
    num.shuffle <- 1000
    res <- sapply(seq_len(num.shuffle), function(i){
        dtt.new <- copy(dtt)
        dtt.new[, miR := sample(miR)]
        x <- dtt.new[tar == 0, miR]
        return('A' %in% x && 'B' %in% x)
    })
    sum(res)/num.shuffle
}

f.exp <- function(x) {(4*x^2 + 6*x + 2) / (9*x^2 + 9*x + 2)}
f.marco <- function(x){4*sqrt(3)*pi / x * (1/3)^(3*x + 3)}

ns <- c(seq(1, 10), seq(20, 100, 10))


set.seed(123)
y.simu <- sapply(ns, simu)
y.exp <- sapply(ns, f.exp)
y.marco <- sapply(ns, f.marco)

pdf('plots/Figure3_simulation.pdf', width = 5, height = 5)
plot(ns, y.simu, ylim = c(0, 1), xlim = c(1, 100), pch = 19,
    xlab = expression(italic(n)), ylab = expression(italic(P)))
curve(f.exp, from = 1, to = 100, add = TRUE, col = 2, lwd = 2)
curve(f.marco, from = 1, to = 100, add = TRUE, col = 4, lwd = 2)
legend(40, 0.3, legend = 
    expression(italic(P)(n) == over(4*sqrt(3)*pi, n)*(over(1,3))^3(n+1)),
    lty = 1, col = 4, bty = 'n', lwd = 2)
legend(40, 0.8, legend =
    expression(italic(P)(n) == over(4*n^2 + 6*n + 2, 9*n^2 + 9*n + 2)),
    lty = 1, col = 2, bty = 'n', lwd = 2)
legend('topleft', legend = 'Simulation', pch = 19, bty = 'n')
dev.off()
```
````
