## Supplementary codes for "Response to “No evidence of functional co-adaptation between clustered microRNAs”": Figure4.Rmd

````
---
title: ""
output: html_notebook
---

###  Figure 4. The distribution of the fraction of common targets for a pair of miRNAs that are from the same cluster (Clustered) or not (Non-clustered).
Paired miRNAs with a seed similarity less
than or equal to six nucleotides were compared separately with the Wilcoxon rank-sum tests.


#### Loading libraries and datasets
```{r}
library(data.table)
library(stringdist)
library(ggplot2)
mytheme <- theme_classic(base_size = 12) + theme(
    axis.text = element_text(color = 'black'),
    strip.background = element_blank(),
    plot.title = element_text(hjust = 0.5),
    plot.subtitle = element_text(hjust = 0.5))

## processed expresion data from Marco (https://doi.org/10.6084/m9.figshare.6165722)
mirna.exp <- fread('data/hsa_miR_binary.tab')
gene.exp <- fread('data/hsa_ensembl_binary.tab')
gene.exp <- gene.exp[!grepl('^__', gene)]

gene.exp.flat <- melt(
    gene.exp, id.vars = 'gene', variable.name = 'tissue', value.name = 'status')
mirna.exp.flat <- melt(
    mirna.exp, id.vars = 'miR', variable.name = 'tissue', value.name = 'status')

# co-expressed gene ~ miRNA pairs
coexp.pair <- merge(
    gene.exp.flat[status > 0, !'status'], mirna.exp.flat[status > 0, !'status'],
    by = 'tissue', allow.cartesian = TRUE)
coexp.pair <- unique(coexp.pair[, !'tissue'])

# miRNA and targets prepared by Marco (https://doi.org/10.6084/m9.figshare.6165722)
mircluster <- fread('data/hsa_miR_clusters.tab', header = FALSE, sep = '\t')
mirfam <- fread('zcat data/miR_Family_mammals_conserved.txt.gz')
mirtar <- fread('zcat data/TargetScan_conserved.txt.gz')
mir2pre <- fread('data/hsa_miR_to_precursor.tab', header = FALSE)

setnames(mircluster, 'V1', 'cluster')
mircluster2 <- mircluster[, .(pre = strsplit(cluster, ';', fixed = TRUE)[[1]]), by = .(cluster)]

mirfam <- mirfam[order(-V7, V4)]
mirfam[, miR := sub('\\.[12]$', '', V4)]

mirtar[, gene := substr(V2, 1, 15)]
mirtar[mirfam, seed := i.V2, on = .(V1)]
mirtar <- unique(mirtar[, .(gene, seed)])

# miRNA, precursor, cluster table
setnames(mir2pre, c('miR', 'pre'))
mir2pre[mircluster2, cluster := i.cluster, on = 'pre']

mircon <- mirfam[grepl('MIMAT', V7), .(miR, family = V1, seed = V2, seq = V5)]
mircon <- merge(mircon, mir2pre, by = 'miR')

head(mircon)
```

#### Analysis
1. Get coexpressed miRNA-target pairs.
```{r}
coexp.pair.fine <- merge(coexp.pair, unique(mircon[, .(miR, seed, pre, cluster)]), by = 'miR')
coexp.pair.fine[mirtar, fzb := 1, on = .(gene, seed)]
coexp.pair.fine <- coexp.pair.fine[!is.na(fzb)]
coexp.pair.fine[, fzb := NULL]

rm(gene.exp, gene.exp.flat, coexp.pair, mirna.exp, mirna.exp.flat)
head(coexp.pair.fine)
```

2. Get common targets of two different miRNAs.
Only pairs of miRNAs that have different seed and from different precursor are considered.
```{r}
common.target <- merge(coexp.pair.fine, coexp.pair.fine, by = 'gene', allow.cartesian = TRUE)
common.target <- common.target[miR.x < miR.y & seed.x != seed.y & pre.x != pre.y]
```

3. Calculate fraction of common targets using harmonic mean of total targets. (Marco used `pmin(total.x, total.y)` to get common targets).
```{r}
target.permir <- coexp.pair.fine[, .(tpm = uniqueN(gene)), by = .(miR)]
dtt <- common.target[, .(common = .N), by = .(miR.x, miR.y, cluster.x, cluster.y, seed.x, seed.y)]
dtt[, sameclust := ifelse(cluster.x == cluster.y, 'Clustered', 'Non-clustered')]
dtt[target.permir, total.x := i.tpm, on = .(miR.x = miR)]
dtt[target.permir, total.y := i.tpm, on = .(miR.y = miR)]
dtt[, total.harm := 2/(1/total.x + 1/total.y)]
dtt[, prob := common / total.harm]
```

4. Stratify by the number of identical nucleotides in seeds
```{r}
dtt[, nuc.ident := 7 - stringdist(seed.x, seed.y)]
dtt2 <- unique(dtt, by = c('miR.x', 'miR.y', 'sameclust'))
```

5. Compare fraction of comments of clustered miRNAs to non-clustered miRNAs (seed similarity < 6):
```{r}
dtt2[nuc.ident < 6,
    wilcox.test(prob[cluster.x == cluster.y], prob[cluster.x != cluster.y])]  # p-value = 0.008054
```

6. Compare fraction of comments of clustered miRNAs to non-clustered miRNAs (seed similarity == 6):
```{r}
dtt2[nuc.ident == 6,
    wilcox.test(prob[cluster.x == cluster.y], prob[cluster.x != cluster.y])]  # p-value = 0.2596
```

7. Produce Figure 4 in biorxiv manuscript:
```{r}
# titles for figure panels
dtt2[, group := ifelse(nuc.ident < 6, 'Seed similarity < 6\n(P=0.008)', 'Seed similarity = 6\n(P=0.26)')]

ggplot(dtt2, aes(x = sameclust, y = prob)) +
    geom_jitter(size = 0.3, width = 0.35) +
    geom_boxplot(aes(color = sameclust), width = 0.8,
        outlier.shape = NA, fill = alpha('#000000', 0), show.legend = FALSE) +
    scale_color_brewer(palette = 'Set1') +
    facet_wrap(~ group) +
    mytheme +
    labs(x = NULL, y = 'Fraction of common targets')
ggsave('plots/Figure4_fraction_of_common_targets_pct.pdf', width = 6, height = 3)

```
````
