## Supplementary codes for "Response to “No evidence of functional co-adaptation between clustered microRNAs”": Figure5.Rmd

````
---
title: ""
output: html_notebook
---

### Figure 5. Permutation analysis of common target genes of clustered miRNAs.

#### 01. Prepare targets with weighted cumulative context score < -0.3
```{bash}
# wget -P data http://www.targetscan.org/vert_71/vert_71_data_download/Summary_Counts.all_predictions.txt.zip
# wget -P data http://www.targetscan.org/vert_71/vert_71_data_download/Gene_info.txt.zip
```

```{r}
suppressPackageStartupMessages(library(Biostrings))
library(data.table)
target.scan <- fread('unzip -c data/Summary_Counts.all_predictions.txt.zip')
setnames(target.scan, make.names(names(target.scan)))

gene.info <- fread('unzip -c data/Gene_info.txt.zip')
setnames(gene.info, make.names(names(gene.info)))

target.scan <- target.scan[Species.ID == 9606,
    .(Transcript.ID, miRNA.family, wcc = Cumulative.weighted.context...score)]
target.scan[, wcc := as.numeric(wcc)]
target.scan <- target.scan[!is.na(wcc)]
target.scan <- target.scan[wcc < -0.3]

target.scan[gene.info, gene:= i.Gene.ID, on = .(Transcript.ID )]
target.scan <- unique(target.scan[, .(gene, seed = miRNA.family)])
target.scan <- target.scan[gene != 'CDR1as']
fwrite(target.scan, file = 'data/TargetScan_wccmd3.txt')

#' get seed and mature sequences of all miRs
hsa.mature <- readDNAStringSet('data/hsa_mature.fa.gz')
hsa.mature <- as(hsa.mature, 'RNAStringSet')

hsa.mature.dt <- data.table(
    miR = names(hsa.mature),
    seq = as.character(hsa.mature),
    seed = as.character(substring(hsa.mature, 2, 8))
)
fwrite(hsa.mature.dt, file = 'data/hsa_mature_seed.tab')
```

#### 02. Panel A
```{r}
#######################################################################################################
# use all miRNAs and all targets with WCC < -0.3
#######################################################################################################
library(parallel)
library(data.table)


FindOverlap <- function(x){
    if(length(x) < 2){
        return(rep(0, length(x)))
    }
    x <- strsplit(x, ';')
    res <- rep(0, length(x))
    for(i in 2:length(x)){
        for(j in 1:(i-1)){
            if(all(x[[i]] %in% x[[j]])){
                res[i] <- j
                break
            }
        }
    }
    return(res)
}

## coexpression informaiton ##############################################################################
mirna.exp <- fread('data/hsa_miR_binary.tab')
gene.exp <- fread('data/hsa_ensembl_binary.tab')
gene.exp <- gene.exp[!grepl('^__', gene)]

gene.exp.flat <- melt(
    gene.exp, id.vars = 'gene', variable.name = 'tissue', value.name = 'status')
mirna.exp.flat <- melt(
    mirna.exp, id.vars = 'miR', variable.name = 'tissue', value.name = 'status')

coexp.pair <- merge(
    gene.exp.flat[status > 0, !'status'], mirna.exp.flat[status > 0, !'status'],
    by = 'tissue', allow.cartesian = TRUE)

coexp.pair <- unique(coexp.pair[, !'tissue'])

## miRNA clusters ########################################################################################
mircluster <- fread('data/hsa_miR_clusters.tab', header = FALSE, sep = '\t')
setnames(mircluster, 'V1', 'cluster')
mircluster2 <- mircluster[, .(pre = strsplit(cluster, ';', fixed = TRUE)[[1]]), by = .(cluster)]

mir2pre <- fread('data/hsa_miR_to_precursor.tab', header = FALSE)
setnames(mir2pre, c('miR', 'pre'))

mirseq <- fread('data/hsa_mature_seed.tab')
mirinfo <- merge(merge(mirseq, mir2pre, by = 'miR'), mircluster2, by = 'pre')

## observed number of genes target by two or more seeds from the same family ###############################
#' target with WCC < -0.3
mirtar <- fread('zcat data/TargetScan_wccmd3.txt.gz')
mirtar[, gene := substr(gene, 1, 15)]

# gene ~ miR ~ seed [gene ~ seed should be in mirtar (fzb: feasible))]
coexp.pair.fine <- merge(coexp.pair, unique(mirinfo[, .(miR, seed)]), by = 'miR')
coexp.pair.fine[mirtar, fzb := 1, on = .(gene, seed)]
coexp.pair.fine <- coexp.pair.fine[!is.na(fzb)]
coexp.pair.fine[, fzb := NULL]

rm(gene.exp, gene.exp.flat, coexp.pair, mirna.exp, mirna.exp.flat)

# get miRNA cluster
gene.mirclust <- merge(
    coexp.pair.fine, unique(mirinfo[, .(miR, cluster, pre)]),
    by = 'miR', allow.cartesian=TRUE)

gene.mirclust[, uniq.seeds := paste(unique(seed), collapse = ';'), by = .(cluster)]

# summary
dtt <- gene.mirclust[, .(
    num_seed = uniqueN(seed), num_pre = uniqueN(pre), seeds = uniq.seeds[1]),
    by = .(gene, cluster)]

dtt2 <- dtt[num_seed > 1 & num_pre > 1][, .(num_gene = uniqueN(gene), seeds = seeds[1]), by = .(cluster)]
dtt2 <- dtt2[order(-nchar(seeds), seeds)]
dtt2[, fov := FindOverlap(seeds)]
obs <- dtt2[fov == 0, sum(num_gene)]


# shuffling miRNA loci #####################################################################################
## Uncommnet of the following block to run simulation

# runs <- 10000
# RNGkind("L'Ecuyer-CMRG")
# set.seed(123)
# marco.loci.shuffle <- mclapply(seq_len(runs), function(i){
#     # shuffle loci
#     mirclust.rand <- mircluster2[, .(cluster.obs = cluster, pre)]
#     mirclust.rand[, pre := sample(pre)]
#     mirclust.rand[, cluster := paste0(pre, collapse = ';'), by = cluster.obs]
#     mirclust.rand[, cluster.obs := NULL]
#
#     mirinfo.rand <- merge(merge(mirseq, mir2pre, by = 'miR'), mirclust.rand, by = 'pre')
#
#     # get common targets
#     gene.mirclust.rand <- merge(
#         coexp.pair.fine, unique(mirinfo.rand[, .(miR, cluster, pre)]),
#         by = 'miR', allow.cartesian=TRUE)
#
#     gene.mirclust.rand[, uniq.seeds := paste(unique(seed), collapse = ';'), by = .(cluster)]
#
#     # summary
#     dtt <- gene.mirclust.rand[, .(
#         num_seed = uniqueN(seed), num_pre = uniqueN(pre), seeds = first(uniq.seeds)),
#         by = .(gene, cluster)]
#
#     dtt2 <- dtt[num_seed > 1 & num_pre > 1]
#     if(nrow(dtt2) == 0){
#         return(0)
#     }
#     dtt2 <- dtt2[, .(num_gene = uniqueN(gene), seeds = seeds[1]), by = .(cluster)]
#     dtt2 <- dtt2[order(-nchar(seeds), seeds)]
#     dtt2[, fov := FindOverlap(seeds)]
#     dtt2[fov == 0, sum(num_gene)]
# }, mc.cores = 20)
# saveRDS(marco.loci.shuffle, file = 'results/FixCoexp_AllMirWCC.rds')

marco.loci.shuffle <- readRDS('results/FixCoexp_AllMirWCC.rds')

simu <- unlist(marco.loci.shuffle)
epvalue <- (sum(simu >= obs) + 1) / (length(simu) + 1)

pdf('plots/Figure5A_FixCoexp_AllMirWCC.pdf', width = 5, height = 4.5)
hist(simu, breaks = 80, xlab = 'Common target genes',
    main = "WCC < -0.3", col = 'grey')
abline(v = obs, col = 'red', lwd = 2)
legend('topright', bty = 'n',
    legend = sprintf("Obs=%d\nP=%.4f", obs, epvalue))
dev.off()
```

#### 03. Panel B
```{r}
#######################################################################################################
#' analysis after excluding mir183~mir182 cluster
#' use all miRNAs and all targets with WCC < -0.3
#' filtering creteria as in Marco: cat datasets/hsa_miR_clusters.tab | grep -v -e'hsa-mir-183' > temp_no183
#######################################################################################################
library(data.table)
library(parallel)

rm(list = ls(all.names = TRUE))

FindOverlap <- function(x){
    if(length(x) < 2){
        return(rep(0, length(x)))
    }
    x <- strsplit(x, ';')
    res <- rep(0, length(x))
    for(i in 2:length(x)){
        for(j in 1:(i-1)){
            if(all(x[[i]] %in% x[[j]])){
                res[i] <- j
                break
            }
        }
    }
    return(res)
}

## coexpression informaiton ##############################################################################
mirna.exp <- fread('data/hsa_miR_binary.tab')
gene.exp <- fread('data/hsa_ensembl_binary.tab')
gene.exp <- gene.exp[!grepl('^__', gene)]

gene.exp.flat <- melt(
    gene.exp, id.vars = 'gene', variable.name = 'tissue', value.name = 'status')
mirna.exp.flat <- melt(
    mirna.exp, id.vars = 'miR', variable.name = 'tissue', value.name = 'status')

coexp.pair <- merge(
    gene.exp.flat[status > 0, !'status'], mirna.exp.flat[status > 0, !'status'],
    by = 'tissue', allow.cartesian = TRUE)

coexp.pair <- unique(coexp.pair[, !'tissue'])

## miRNA clusters ########################################################################################
mircluster <- fread(
    cmd = 'grep -v -e "hsa-mir-183" data/hsa_miR_clusters.tab', header = FALSE, sep = '\t')
setnames(mircluster, 'V1', 'cluster')
mircluster2 <- mircluster[, .(pre = strsplit(cluster, ';', fixed = TRUE)[[1]]), by = .(cluster)]

mir2pre <- fread('data/hsa_miR_to_precursor.tab', header = FALSE)
setnames(mir2pre, c('miR', 'pre'))

mirseq <- fread('data/hsa_mature_seed.tab')
mirinfo <- merge(merge(mirseq, mir2pre, by = 'miR'), mircluster2, by = 'pre')

## observed number of genes target by two or more seeds from the same family #############################
#' target with WCC < -0.3
mirtar <- fread('zcat data/TargetScan_wccmd3.txt.gz')
mirtar[, gene := substr(gene, 1, 15)]

# gene ~ miR ~ seed [gene ~ seed should be in mirtar (fzb: feasible))]
coexp.pair.fine <- merge(coexp.pair, unique(mirinfo[, .(miR, seed)]), by = 'miR')
coexp.pair.fine[mirtar, fzb := 1, on = .(gene, seed)]
coexp.pair.fine <- coexp.pair.fine[!is.na(fzb)]
coexp.pair.fine[, fzb := NULL]

rm(gene.exp, gene.exp.flat, coexp.pair, mirna.exp, mirna.exp.flat)

# get miRNA cluster
gene.mirclust <- merge(
    coexp.pair.fine, unique(mirinfo[, .(miR, cluster, pre)]),
    by = 'miR', allow.cartesian=TRUE)

gene.mirclust[, uniq.seeds := paste(unique(seed), collapse = ';'), by = .(cluster)]

# summary
dtt <- gene.mirclust[, .(
    num_seed = uniqueN(seed), num_pre = uniqueN(pre), seeds = first(uniq.seeds)),
    by = .(gene, cluster)]

dtt2 <- dtt[num_seed > 1 & num_pre > 1][, .(num_gene = uniqueN(gene), seeds = seeds[1]), by = .(cluster)]
dtt2 <- dtt2[order(-nchar(seeds), seeds)]
dtt2[, fov := FindOverlap(seeds)]
obs <- dtt2[fov == 0, sum(num_gene)]


# shuffling miRNA loci ###################################################################################
# runs <- 10000
# RNGkind("L'Ecuyer-CMRG")
# set.seed(123)
# marco.loci.shuffle <- mclapply(seq_len(runs), function(i){
#     # shuffle loci
#     mirclust.rand <- mircluster2[, .(cluster.obs = cluster, pre)]
#     mirclust.rand[, pre := sample(pre)]
#     mirclust.rand[, cluster := paste0(pre, collapse = ';'), by = cluster.obs]
#     mirclust.rand[, cluster.obs := NULL]
#
#     mirinfo.rand <- merge(merge(mirseq, mir2pre, by = 'miR'), mirclust.rand, by = 'pre')
#
#     # get common targets
#     gene.mirclust.rand <- merge(
#         coexp.pair.fine, unique(mirinfo.rand[, .(miR, cluster, pre)]),
#         by = 'miR', allow.cartesian=TRUE)
#
#     gene.mirclust.rand[, uniq.seeds := paste(unique(seed), collapse = ';'), by = .(cluster)]
#
#     # summary
#     dtt <- gene.mirclust.rand[, .(
#         num_seed = uniqueN(seed), num_pre = uniqueN(pre), seeds = first(uniq.seeds)),
#         by = .(gene, cluster)]
#
#     dtt2 <- dtt[num_seed > 1 & num_pre > 1]
#     if(nrow(dtt2) == 0){
#         return(0)
#     }
#     dtt2 <- dtt2[, .(num_gene = uniqueN(gene), seeds = seeds[1]), by = .(cluster)]
#     dtt2 <- dtt2[order(-nchar(seeds), seeds)]
#     dtt2[, fov := FindOverlap(seeds)]
#     dtt2[fov == 0, sum(num_gene)]
# }, mc.cores = 20)
#
# saveRDS(marco.loci.shuffle, file = 'results/FixCoexp_AllMirWCC_no183.rds')

marco.loci.shuffle <- readRDS('results/FixCoexp_AllMirWCC_no183.rds')

simu <- unlist(marco.loci.shuffle)
epvalue <- (sum(simu >= obs) + 1) / (length(simu) + 1)

pdf('plots/Figure5B_FixCoexp_AllMirWCC_no183.pdf', width = 5, height = 4.5)
hist(simu, breaks = 80, xlab = 'Common target genes',
    main = "WCC < -0.3 (Excluding mir-183~182)", col = 'grey')
abline(v = obs, col = 'red', lwd = 2)
legend('topright', bty = 'n',
    legend = sprintf("Obs=%d\nP=%.4f", obs, epvalue))
dev.off()
```

#### 04. Panel C
```{r}
#######################################################################################################
#' use all miRNAs and all targets with WCC < -0.3
#######################################################################################################
library(data.table)
library(parallel)
rm(list = ls(all.names = TRUE))

FindOverlap <- function(x){
    if(length(x) < 2){
        return(rep(0, length(x)))
    }
    x <- strsplit(x, ';')
    res <- rep(0, length(x))
    for(i in 2:length(x)){
        for(j in 1:(i-1)){
            if(all(x[[i]] %in% x[[j]])){
                res[i] <- j
                break
            }
        }
    }
    return(res)
}

## coexpression informaiton ##############################################################################
mirna.exp <- fread('data/hsa_miR_binary.tab')
gene.exp <- fread('data/hsa_ensembl_binary.tab')
gene.exp <- gene.exp[!grepl('^__', gene)]

gene.exp.flat <- melt(
    gene.exp, id.vars = 'gene', variable.name = 'tissue', value.name = 'status')
mirna.exp.flat <- melt(
    mirna.exp, id.vars = 'miR', variable.name = 'tissue', value.name = 'status')

coexp.pair <- merge(
    gene.exp.flat[status > 0, !'status'], mirna.exp.flat[status > 0, !'status'],
    by = 'tissue', allow.cartesian = TRUE)

coexp.pair <- unique(coexp.pair[, !'tissue'])

## miRNA clusters ########################################################################################
mircluster <- fread('data/hsa_miR_clusters.tab', header = FALSE, sep = '\t')
setnames(mircluster, 'V1', 'cluster')
mircluster2 <- mircluster[, .(pre = strsplit(cluster, ';', fixed = TRUE)[[1]]), by = .(cluster)]

mir2pre <- fread('data/hsa_miR_to_precursor.tab', header = FALSE)
setnames(mir2pre, c('miR', 'pre'))

mirseq <- fread('data/hsa_mature_seed.tab')
mirinfo <- merge(merge(mirseq, mir2pre, by = 'miR'), mircluster2, by = 'pre')

## observed number of genes target by two or more seeds from the same family ###############################
#' target with WCC < -0.3
mirtar <- fread('zcat data/TargetScan_wccmd3.txt.gz')
mirtar[, gene := substr(gene, 1, 15)]

# gene ~ miR ~ seed [gene ~ seed should be in mirtar (fzb: feasible))]
coexp.pair.fine <- merge(coexp.pair, unique(mirinfo[, .(miR, seed)]), by = 'miR')
coexp.pair.fine[mirtar, fzb := 1, on = .(gene, seed)]
coexp.pair.fine <- coexp.pair.fine[!is.na(fzb)]
coexp.pair.fine[, fzb := NULL]
coexp.pair.fine <- unique(coexp.pair.fine[, .(gene, seed)])

rm(gene.exp, gene.exp.flat, coexp.pair, mirna.exp, mirna.exp.flat)

# shuffling miRNA loci #####################################################################################
# runs <- 10000
# RNGkind("L'Ecuyer-CMRG")
# set.seed(123)
# coexp.pair.shuffle <- mclapply(seq_len(runs), function(i){
#     # shuffle coexpressed miR-target pairs
#     x <- sample.int(nrow(coexp.pair.fine))
#     coexp.pair.rand <- coexp.pair.fine[, .(gene, seed = seed[x])]
#
#     # get common targets
#     gene.mirclust <- merge(
#         coexp.pair.rand, unique(mirinfo[, .(seed, cluster, pre)]),
#         by = 'seed', allow.cartesian=TRUE)
#
#     gene.mirclust[, uniq.seeds := paste(unique(seed), collapse = ';'), by = .(cluster)]
#
#     # summary
#     dtt <- gene.mirclust[, .(
#         num_seed = uniqueN(seed), num_pre = uniqueN(pre), seeds = first(uniq.seeds)),
#         by = .(gene, cluster)]
#
#     obs <- dtt[num_seed > 1, uniqueN(gene)]
# }, mc.cores = 10)
#
# saveRDS(coexp.pair.shuffle, file = 'results/ShufflePairOldMeth_FixCoexp_AllMirWCC.rds')

coexp.pair.shuffle <- readRDS('results/ShufflePairOldMeth_FixCoexp_AllMirWCC.rds')

obs <- 2081
simu <- unlist(coexp.pair.shuffle)
epvalue <- (sum(simu >= obs) + 1) / (length(simu) + 1)

pdf('plots/Figure5C_ShufflePairOldMeth_FixCoexp_AllMirWCC.pdf', width = 5, height = 4.5)
hist(simu, breaks = seq(1620, 2200, 10), xlab = 'Common target genes',
    main = "WCC < -0.3", col = 'grey')
abline(v = obs, col = 'red', lwd = 2)
legend('topright', bty = 'n',
    legend = sprintf("Observed=%d\nP=%.4f", obs, epvalue))
dev.off()

```

#### 05. Panel D
```{r}
## general notes ######################################################################################
#' use all miRNAs and all targets with WCC < -0.3 after excluding mir-183~182 cluster
#######################################################################################################
library(data.table)
library(parallel)
rm(list = ls(all.names = TRUE))

FindOverlap <- function(x){
    if(length(x) < 2){
        return(rep(0, length(x)))
    }
    x <- strsplit(x, ';')
    res <- rep(0, length(x))
    for(i in 2:length(x)){
        for(j in 1:(i-1)){
            if(all(x[[i]] %in% x[[j]])){
                res[i] <- j
                break
            }
        }
    }
    return(res)
}

## coexpression informaiton ##############################################################################
mirna.exp <- fread('data/hsa_miR_binary.tab')
gene.exp <- fread('data/hsa_ensembl_binary.tab')
gene.exp <- gene.exp[!grepl('^__', gene)]

gene.exp.flat <- melt(
    gene.exp, id.vars = 'gene', variable.name = 'tissue', value.name = 'status')
mirna.exp.flat <- melt(
    mirna.exp, id.vars = 'miR', variable.name = 'tissue', value.name = 'status')

coexp.pair <- merge(
    gene.exp.flat[status > 0, !'status'], mirna.exp.flat[status > 0, !'status'],
    by = 'tissue', allow.cartesian = TRUE)

coexp.pair <- unique(coexp.pair[, !'tissue'])

## miRNA clusters ########################################################################################
mircluster <- fread(
    cmd = 'grep -v -e "hsa-mir-183" data/hsa_miR_clusters.tab', header = FALSE, sep = '\t')
setnames(mircluster, 'V1', 'cluster')
mircluster2 <- mircluster[, .(pre = strsplit(cluster, ';', fixed = TRUE)[[1]]), by = .(cluster)]

mir2pre <- fread('data/hsa_miR_to_precursor.tab', header = FALSE)
setnames(mir2pre, c('miR', 'pre'))

mirseq <- fread('data/hsa_mature_seed.tab')
mirinfo <- merge(merge(mirseq, mir2pre, by = 'miR'), mircluster2, by = 'pre')

## observed number of genes target by two or more seeds from the same family ###############################
#' target with WCC < -0.3
mirtar <- fread('zcat data/TargetScan_wccmd3.txt.gz')
mirtar[, gene := substr(gene, 1, 15)]

# gene ~ miR ~ seed [gene ~ seed should be in mirtar (fzb: feasible))]
coexp.pair.fine <- merge(coexp.pair, unique(mirinfo[, .(miR, seed)]), by = 'miR')
coexp.pair.fine[mirtar, fzb := 1, on = .(gene, seed)]
coexp.pair.fine <- coexp.pair.fine[!is.na(fzb)]
coexp.pair.fine[, fzb := NULL]
coexp.pair.fine <- unique(coexp.pair.fine[, .(gene, seed)])

rm(gene.exp, gene.exp.flat, coexp.pair, mirna.exp, mirna.exp.flat)

# shuffling miRNA loci #####################################################################################
# runs <- 10000
#
# RNGkind("L'Ecuyer-CMRG")
# set.seed(123)
# coexp.pair.shuffle <- mclapply(seq_len(runs), function(i){
#     # shuffle coexpressed miR-target pairs
#     x <- sample.int(nrow(coexp.pair.fine))
#     coexp.pair.rand <- coexp.pair.fine[, .(gene, seed = seed[x])]
#
#     # get common targets
#     gene.mirclust <- merge(
#         coexp.pair.rand, unique(mirinfo[, .(seed, cluster, pre)]),
#         by = 'seed', allow.cartesian=TRUE)
#
#     gene.mirclust[, uniq.seeds := paste(unique(seed), collapse = ';'), by = .(cluster)]
#
#     # summary
#     dtt <- gene.mirclust[, .(
#         num_seed = uniqueN(seed), num_pre = uniqueN(pre), seeds = first(uniq.seeds)),
#         by = .(gene, cluster)]
#
#     obs <- dtt[num_seed > 1, uniqueN(gene)]
# }, mc.cores = 10)
#
# saveRDS(coexp.pair.shuffle, file = 'results/ShufflePairOldMeth_FixCoexp_AllMirWCC_no183.rds')

coexp.pair.shuffle <- readRDS('results/ShufflePairOldMeth_FixCoexp_AllMirWCC_no183.rds')

obs <- 1956
simu <- unlist(coexp.pair.shuffle)
epvalue <- (sum(simu >= obs) + 1) / (length(simu) + 1)

pdf('plots/Figure5D_ShufflePairOldMeth_FixCoexp_AllMirWCC_no183.pdf', width = 5, height = 4.5)
hist(simu, breaks = seq(1620, 2000, 10), xlab = 'Common target genes',
    main = "WCC < -0.3 (Excluding mir-183~182)", col = 'grey')
abline(v = obs, col = 'red', lwd = 2)
legend('topright', bty = 'n',
    legend = sprintf("Observed=%d\nP=%.4f", obs, epvalue))
dev.off()

```
````
