## Supplementary figures and images for "Response to “No evidence of functional co-adaptation between clustered microRNAs”"

### Figure1_shuffle_site.pdf

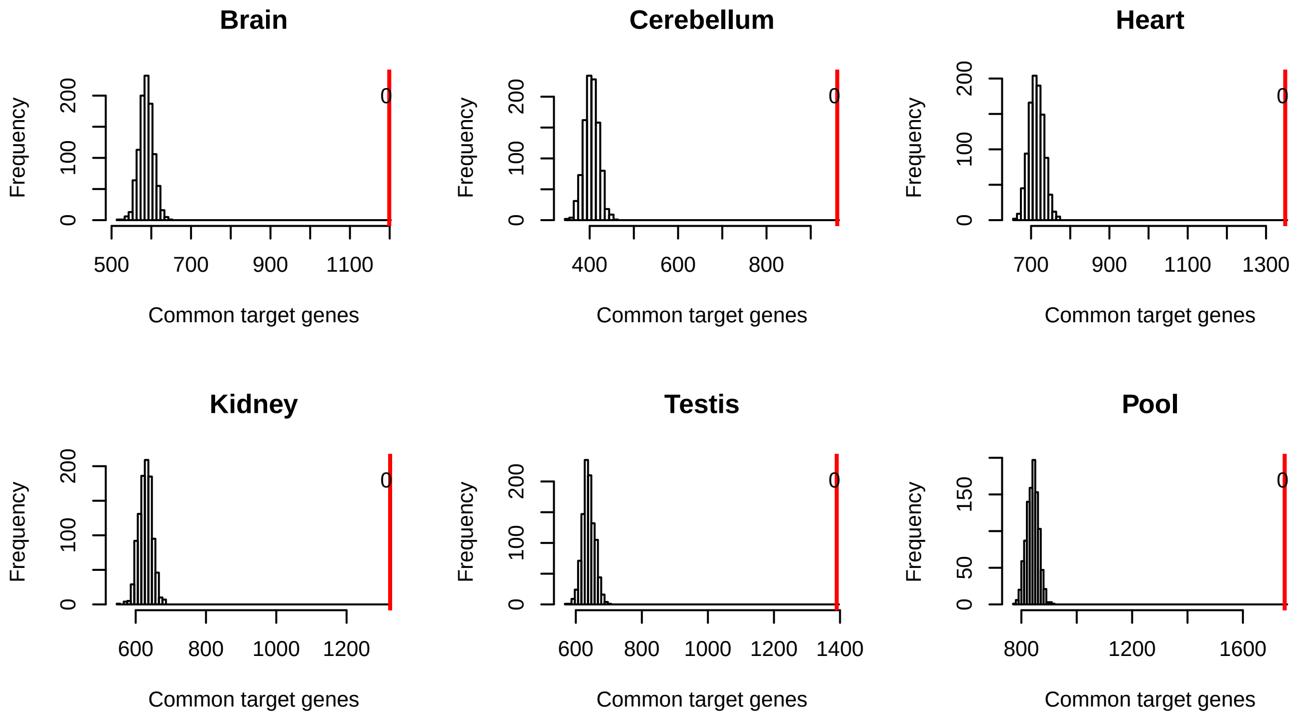

### Figure2_shuffle_site_no183.pdf

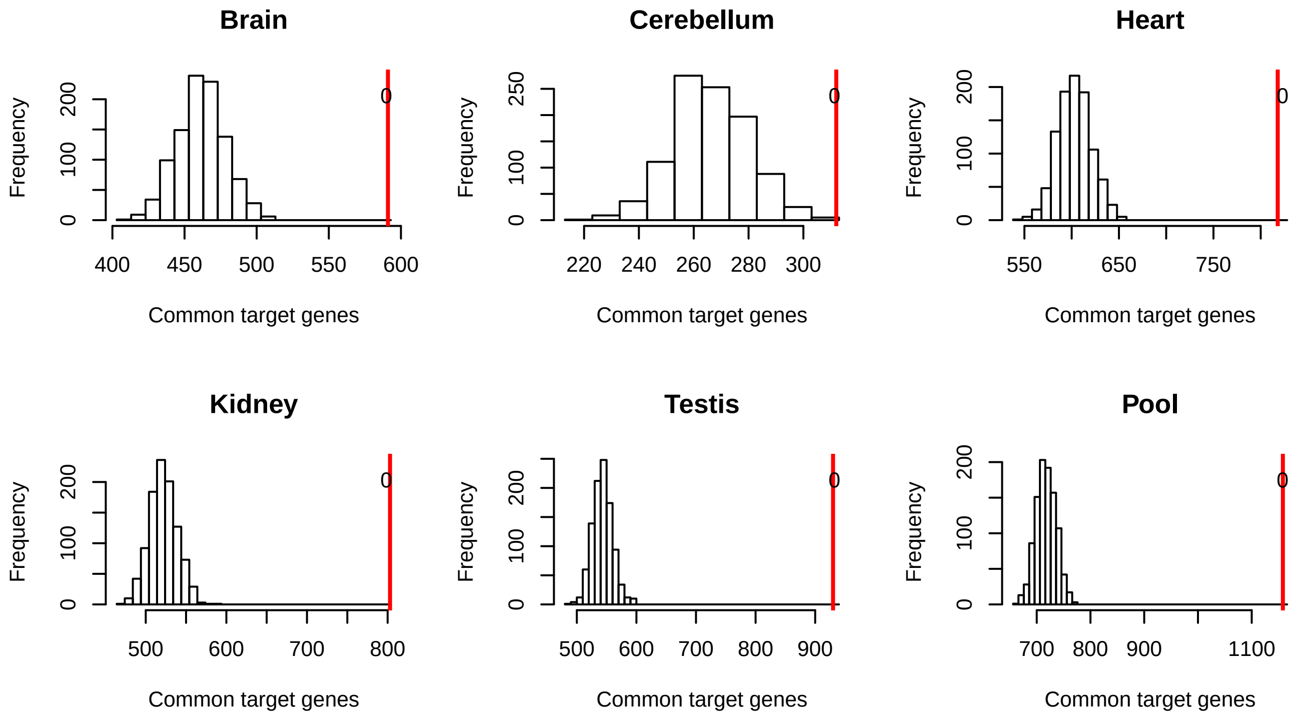

### Figure3_simulation.pdf

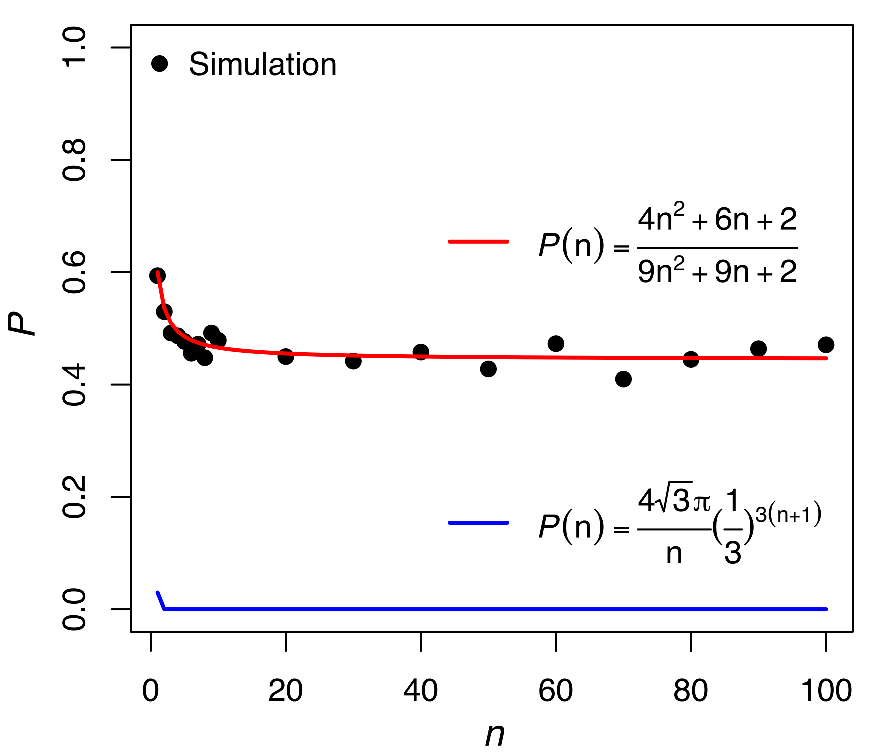

### Figure4_fraction_of_common_targets_pct.pdf

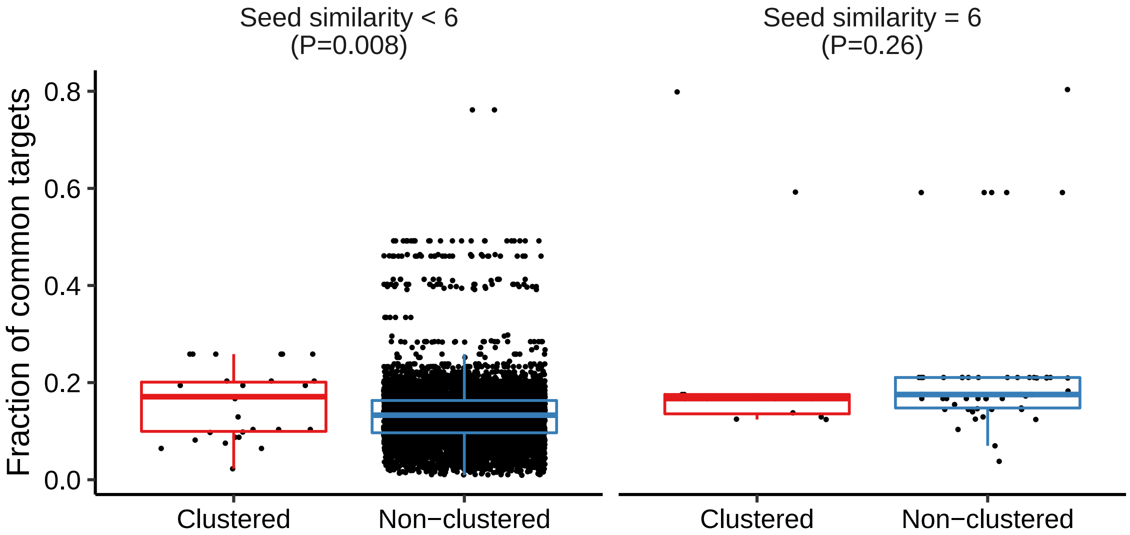

### Figure5A_FixCoexp_AllMirWCC.pdf

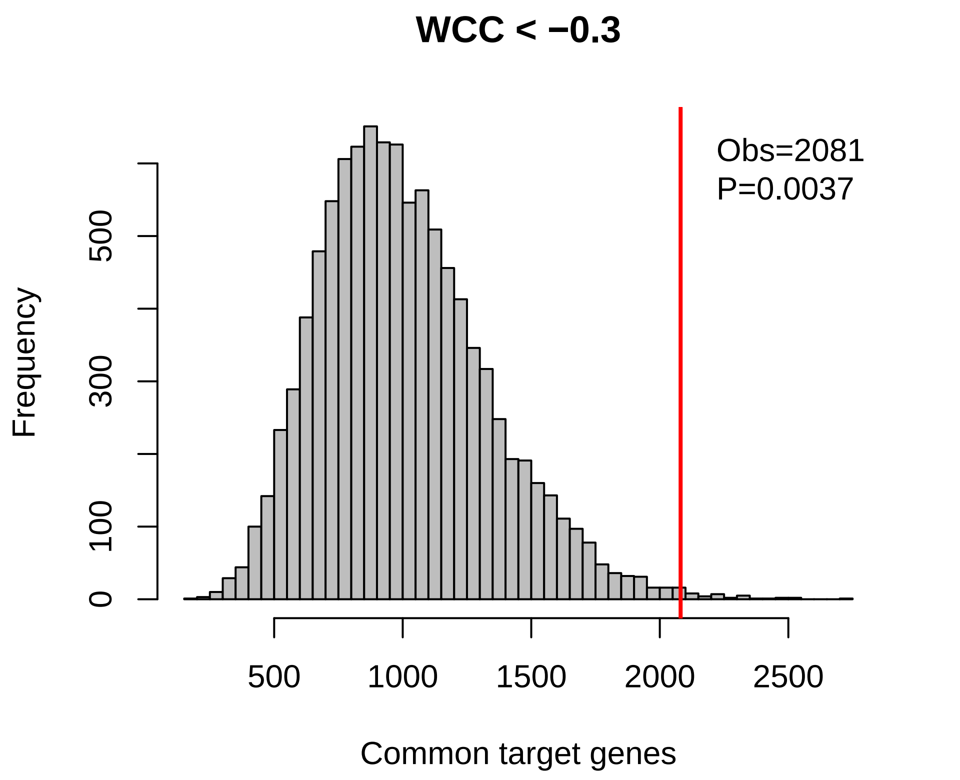

### Figure5B_FixCoexp_AllMirWCC_no183.pdf

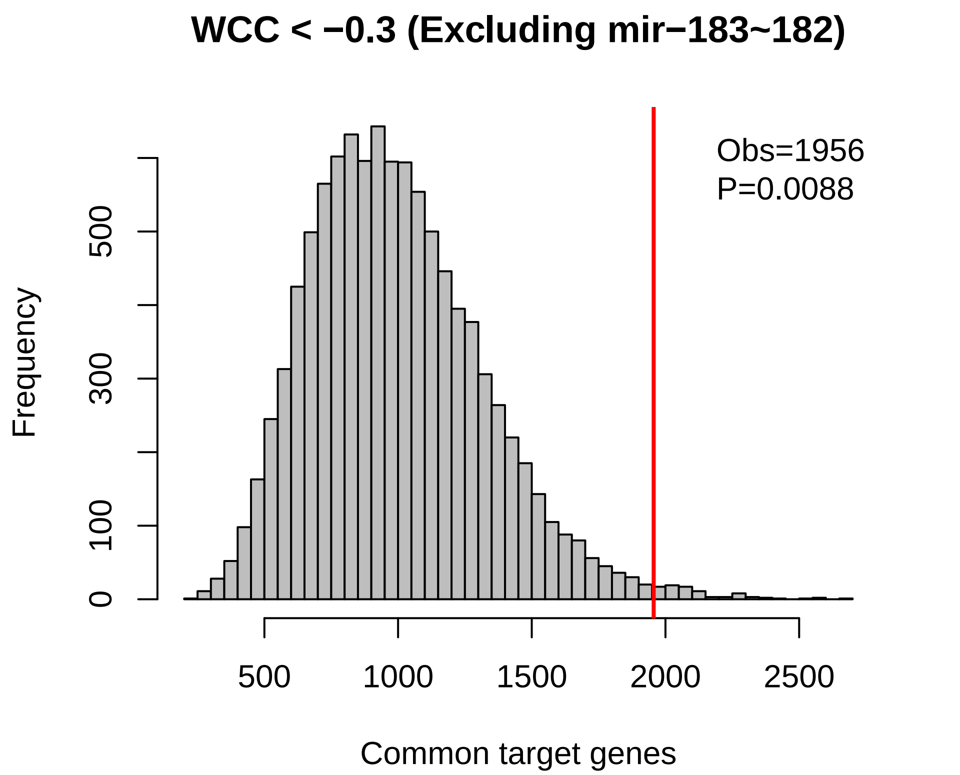

### Figure5C_ShufflePairOldMeth_FixCoexp_AllMirWCC.pdf

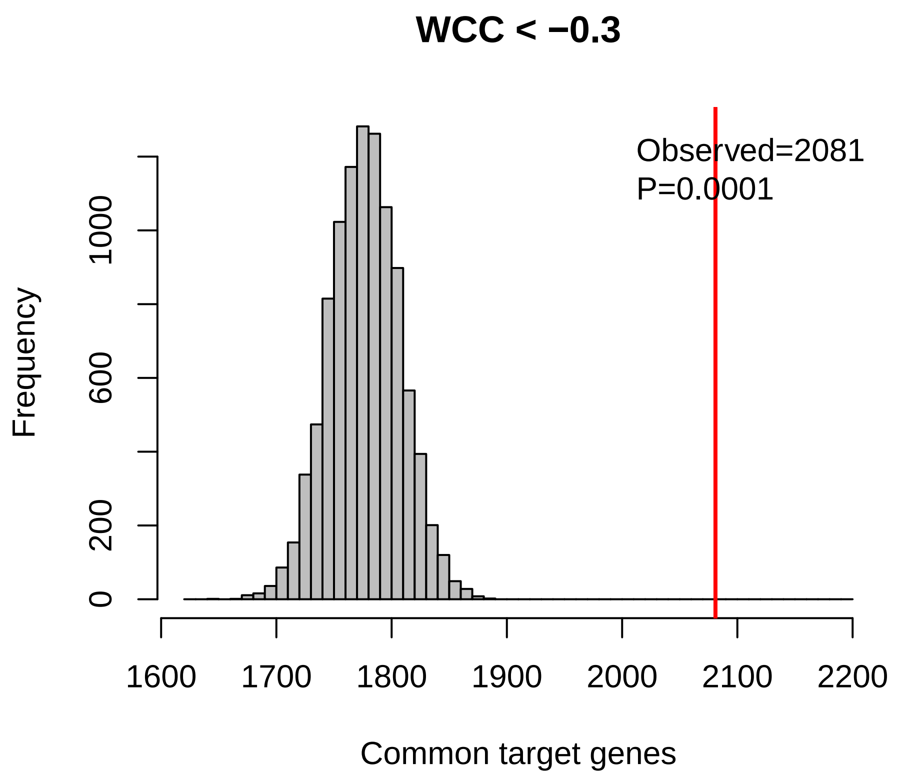

### Figure5D_ShufflePairOldMeth_FixCoexp_AllMirWCC_no183.pdf

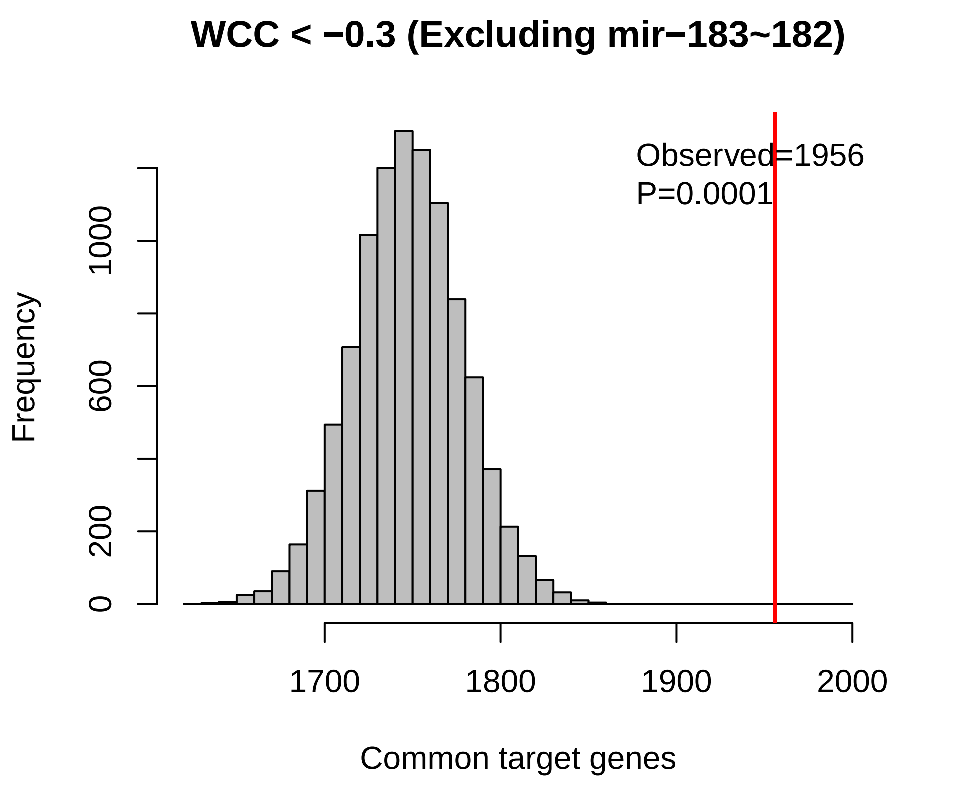

### Figure6_shuffle_site_mir17_92.pdf

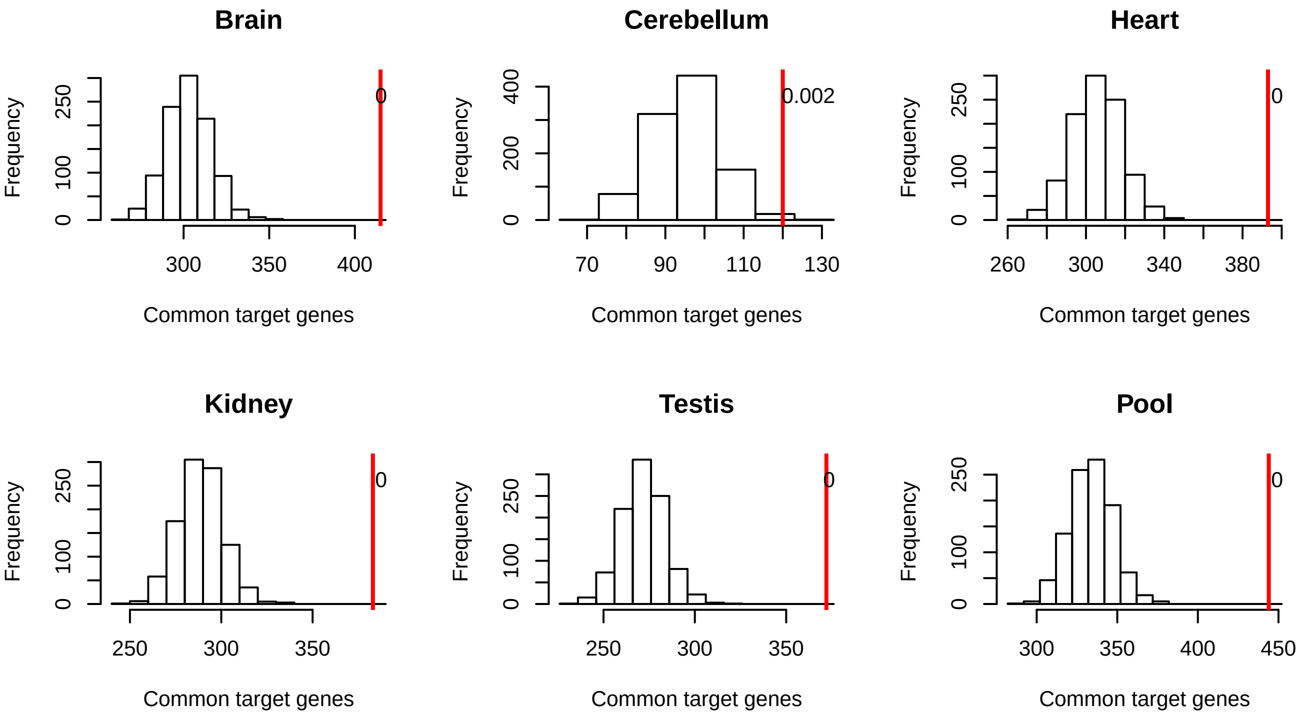
